# Welding PROxAb Shuttles: A modular approach for generating bispecific antibodies via site-specific protein-protein conjugation

**DOI:** 10.1101/2024.03.15.585232

**Authors:** Tanja Lehmann, Hendrik Schneider, Jason Tonillo, Jennifer Schanz, Daniel Schwarz, Christian Schröter, Harald Kolmar, Stefan Hecht, Jan Anderl, Nicolas Rasche, Marcel Rieker, Stephan Dickgiesser

**Affiliations:** Clemens-Schöpf-Institut für Organische Chemie und Biochemie, Technische Universität Darmstadt, 64287 Darmstadt, Germany; Merck KGaA, 64293 Darmstadt, Germany

**Keywords:** PROTAC, antibody, antibody drug conjugate, targeted protein degradation, MTG, bioconjugation

## Abstract

Targeted protein degradation is an innovative therapeutic strategy to selectively eliminate disease-causing proteins. Exemplified by proteolysis-targeting chimeras (PROTACs), it has shown promise in overcoming drug resistance and targeting previously undruggable proteins. However, PROTACs face challenges such as low oral bioavailability and limited selectivity. The recently published PROxAb Shuttle technology offers a solution enabling targeted delivery of PROTACs using antibodies fused with PROTAC-binding domains derived from camelid single-domain antibodies (VHHs). Here, a modular approach to quickly generate PROxAb Shuttles by enzymatically coupling PROTAC-binding VHHs to off-the-shelf antibodies was developed. The resulting conjugates retained their target binding and internalization properties and incubation with BRD4-targeting PROTACs resulted in formation of defined PROxAb-PROTAC complexes. These complexes selectively induced degradation of the BRD4 protein, resulting in cytotoxicity specifically in cells expressing the antibody’s target. The chemoenzymatic approach described here provides a versatile and efficient solution for generating antibody-VHH conjugates for targeted protein degradation applications but could also be used to combine antibody and VHH binders to generate bispecific antibodies for further applications.

## Introduction

Targeted protein degradation (TPD) is a promising new therapeutic strategy that has gained considerable attention in drug discovery. The approach offers an alternative mode of action compared to traditional small molecule inhibitors and antagonists by inducing selective degradation of target proteins. By eliminating disease-causing proteins rather than simply inhibiting their activity, TPD is a promising approach for previously undruggable targets, inducing enhanced responses, and overcoming resistance mechanisms. Among the most prominent targeted degraders are hetero-bifunctional molecules called proteolysis-targeting chimeras (PROTACs) which are composed of two ligands that bind the E3 ligase and the target protein (POI) and are connected by a linker moiety. Simultaneous binding of the E3 ligase and the POI leads to the formation of a ternary complex that induces E3 ligase-mediated polyubiquitination of the POI followed by proteasomal POI degradation via the ubiquitin-proteasome system (UPS).^1^ The PROTAC itself is not degraded during this process and therefore available to recruit the next target molecule in a catalytic mechanism of action.^2^

Unlike traditional inhibitors that require active binding sites to exert their effect, PROTACs can potentially engage target molecules that lack active binding sites and are thus inaccessible to small molecule inhibitors. This increases the likelihood to find suitable ligands especially since low-affinity binding can already be sufficient for ternary complex formation.^3^ Hence, over thousand small molecule PROTACs have been identified to date targeting a wide range of POIs with important biological functions, particularly in cancer cells, including BET-bromodomain proteins^4^, estrogen receptor^5^, androgen receptor^6^ and various kinases.^7, 8, 9^ More than 10 PROTACs have even been advanced to clinical trials already.^10, 11, 12^

While PROTACs hold great promise, they face several challenges for optimal efficacy in clinical applications. They are relatively large molecules (molecular weight >800 Da) and often suffer from low oral bioavailability, cell permeability, and water solubility.^13^ The limited serum stability and cell permeability negatively affect their pharmacokinetics and bioavailability.^14^ Also, selectivity is often limited and can result in off-target effects as E3 ligases deployed for protein degradation have a broad expression spectrum in both tumor and healthy tissues. Therefore, enhancing PROTAC delivery and selectivity is a crucial focus in PROTAC development.^15, 16^ Various approaches are being explored such as encapsulation in lipid nanoparticles for improved cell internalization and lysosomal escape^17^ or fusion to target-selective peptides or chemical groups.^18^ Small-molecule binders, e.g. directed against folate groups, can also serve as ligands for selective PROTAC delivery, as demonstrated by Lui and colleagues.^19^ Notably, antibody-PROTAC conjugates that, similar to antibody-drug conjugates (ADC), combine the target specificity and favorable pharmacokinetic profile of monoclonal antibodies (mAb) with PROTAC-induced protein degradation, lead to enhanced exposure at target cells and potentially reduced off-target effects.^20, 21^ PROTACs frequently lack chemical groups like primary, secondary, or tertiary amines that are typically used for covalent attachment and seamless in situ removal of cleavable linkers for ADCs.^22^ Thus, a key challenge is the complex synthesis of PROTACs, which makes it difficult to incorporate a suitable exit vector or chemical functionalities.

The PROxAb technology is a recently described approach that allows targeted delivery of PROTACs in an ADC-like manner without the need for chemical PROTAC modification.^23^ PROxAb Shuttles are IgG antibodies directed against e.g. tumor-associated antigens fused with variable domains of camelid single-domain antibodies (VHH) that specifically bind the E3 ligase-recruiting subunit of PROTACs and thereby enable formation of non-covalent antibody-PROTAC complexes.^24, 25^ Schneider and colleagues described the discovery of the VHH ’MIC7’, which binds with high affinity and specificity to VH032 – a recruiting domain for the E3 ligase Von Hippel-Lindau tumor suppressor (VHL). As most PROTACs developed to date recruit either of the two E3 ligases VHL and cereblon (CRBN), PROxAb Shuttles are applicable to a broad range of already existing molecules.^26, 27, 28^ Simple incubation of PROTACs with MIC7-based antibody-fusion proteins yielded antibody-PROTAC complexes that induced degradation of PROTAC target protein specifically in cells expressing the antibody target on the cell surface. ^23^ *In vivo* studies demonstrated that complexation positively impacts PROTAC half-life in PK studies and leads PROxAb Shuttle a non-covalent plug-and-play platform for the rapid generation of tumor-targeting antibody-PROTAC conjugates to prolonged anti-tumor effects compared to free PROTAC.

While the PROxAb Shuttle technology eliminates the need for covalent PROTAC modification, it requires individual recombinant production of every antibody-VHH combination. To enhance the flexibility and modularity of the PROxAb Shuttle technology even further, we envisioned biochemical conjugation of PROTAC-binding VHHs to already existing, non-engineered antibodies. This would enable the conversion of any commercially available IgG1-based antibody into a PROxAb shuttle and thereby enable rapid screening of antibody-PROTAC combinations. Among the technologies available for site-specific conjugation of native antibodies, we selected enzymatic coupling via microbial transglutaminase (MTG). MTG catalyzes the formation of isopeptide bonds between glutamine residues and acyl acceptor substrates. Our group has demonstrated previously, that certain MTGs can specifically address the glutamine 295 (Q295; EU numbering) in the heavy chain (HC) of fully glycosylated antibodies without the need for prior insertion of MTG recognition sequences.^29, 30^ Until now, this method has primarily been shown to be effective for attaching small molecules or peptides.^31^ Hence, the objective of this study is to extend and optimize its application for protein-protein conjugation, particularly for the attachment of VHH MIC7 (Figure 1).

**Figure 1.**
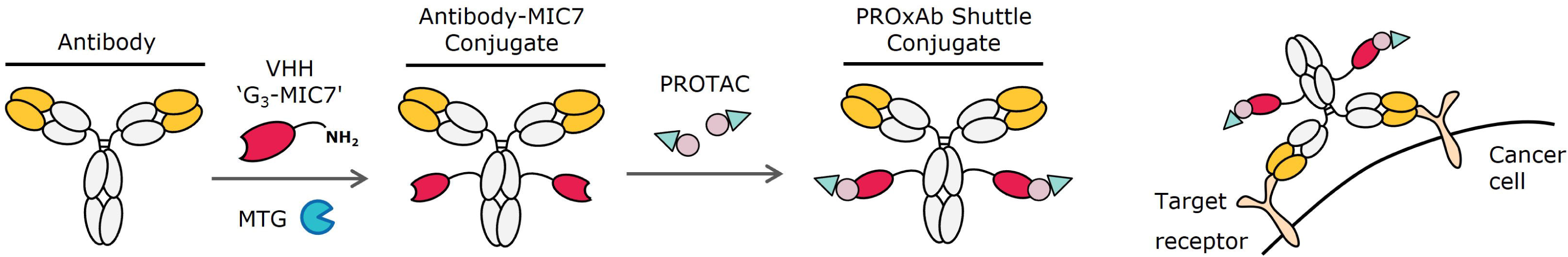
Principle of the modular to generate PROTAC-delivering PROxAb shuttles. Off-the-shelf IgG-based antibodies are site-specifically coupled with VHHs using microbial transglutaminase (MTG). The conjugated PROxAb shuttle as bispecific protein is able to specifically bind PROTACs which is mediated by G_3_-MIC7. Incubation with PROTACs results in formation of stable complexes that induce selective target protein degradation and cytotoxicity exclusively in the target cells.

A protein that is often targeted for PROTAC-induced degradation is the transcriptional regulator bromodomain-containing protein 4 (BRD4) from the BET-family.^22^ Hence, several PROTACs targeting BRD4 already exist and various recruit the E3 ligase VHL.^32^ We selected three of such for the underlying work: GNE987, GNE987P and ARV771.^33^. ^34, 32^ GNE987 and GNE987P are closely related, but while GNE987 utilizes an aliphatic linker, the GNE987P subunits are connected by a hydrophilic polyethylenglycol chain (structure in supplemental section, Figure S1).^34^ This subtle difference leads to longer ternary complex half-life, and increased hydrophobicity and cell permeability of GNE987 compared to GNE987P.^34^

Herein, we present a modular approach for rapid generation of PROTAC-delivering antibodies from off-the-shelf, native mAbs (Figure 1). Therefore, a PROTAC-binding VHH was equipped with an N-terminal triple-glycine motif to allow recognition by MTG^35^ for site-specific conjugation to IgG-based antibodies. The produced conjugates showed the same binding and internalization properties as the parental antibodies. Incubation with PROTACs led to the formation of stable complexes that mediated target protein degradation and cell cytotoxicity selectively to target cells only.

## Results and discussion

### MTG conjugation allows to attach PROTAC-binding VHH to native antibodies

We conjugated PROTAC-binding VHHs to any native, fully glycosylated IgG1 mAb in a single step using enzymatic MTG conjugation targeting Q295 of the mAb heavy chain (Figure 1). Resulting VHH-antibody conjugates function as ‘PROxAb shuttles’ by complexing PROTACs and delivering it to the target the antibody is direct against. To allow for VHH-antibody conjugation, VHH ‘MIC7’ was equipped recombinantly with an N-terminal triple-glycine motif meant to serve as an MTG recognition tag, followed by a glycine-serine spacer. The modified VHH G_3_-MIC7 was expressed in HEK293F cells and purified by affinity chromatography to high monomeric purity of >99.3 % as determined by analytical size exclusion chromatography (SEC) (Figure S2). To establish our solution, we used three IgG1 antibodies as model systems: human epidermal growth factor receptor 2 (HER2) binding Trastuzumab (Ttz; Roche), human epidermal growth factor receptor binding Cetuximab (Ctx, Merck KGaA) as well as digoxigenin-binding αDIG).^36, 37^ Due to the homodimeric architecture of IgGs, site-specific conjugation typically results in an antibody with two VHH molecules attached. 5 mg/ml Ttz were incubated with 10 molar equivalents G_3_-MIC7 and 10 U/mL MTG (produced as described before^30^) for 24 h at 30 °C. Reduced SDS-PAGE analysis revealed a shift of the antibody heavy chain (HC) towards higher molecular weight only in the mix containing all reaction components, indicating successful attachment of the VHH to the HC (Figure 2 (A)).

**Figure 2.**
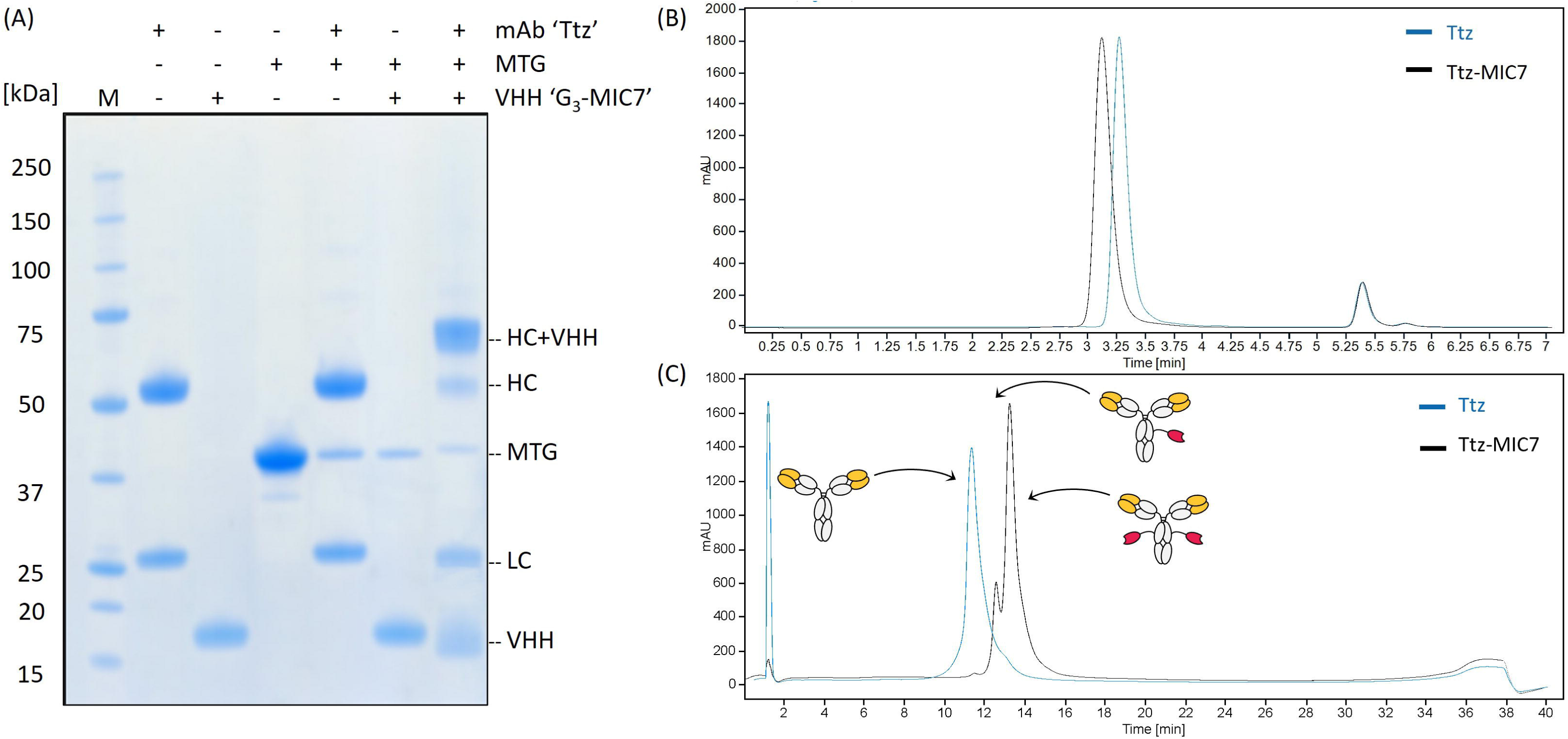
Enzymatic conjugation of VHH ‘G_3_-MIC7’ to Ttz to generate PROxAb shuttle Ttz-MIC7. (A) Reduced SDS-PAGE analysis of individual reactants and reaction mixes. A shift of the Ttz HC but not the LC) band in the final reaction mix indicates successful MTG-driven conjugation of MIC7 to the HC only. (B) Analytical SEC shows a shift towards higher molecular weight in the expected range but does not indicate formation of larger species. (C) HIC analysis of SEC-purified product reveals full conversion of Ttz to two species: a small peak assigned to species with 1 MIC7 attached and a second, major peak representing the anticipated conjugate with 2 MIC7 molecules attached.

SEC analysis of the reaction mix showed an apparent monomeric purity of the product of >97 % suggesting no significant formation of high molecular weight species or non-reacted mAb (Figure S3). In order to confirm site-specific conjugation to the targeted position within the antibody, a variant of trastuzumab with a glutamine to alanine substitution at HC position Q295 was included as control.^30^ Conjugation reactions with this variant did not result in any conjugation products confirming Q295 as the only position targeted by MTG by this approach (Figure S8).

After assessing feasibility of our approach, we sought to produce and purify mAb-MIC7 conjugates for further experiments. A demonstration of chromatographic purification and analysis is shown using Ttz as an example, with the results for all antibodies listed in Table 1. Therefore, 2.0-5.0 mg scale reactions using Ttz, Ctx and αDIG antibody were set up and purified by preparative SEC. Minor amounts of aggregates formed during the reaction (Figure S4 (A)) were removed during purification resulting in conjugates of high purity (Figure 2 (B)) and conjugate yields >72 % (Figure S4 (B)). Hydrophobic interaction chromatography (HIC) (Figure 2 (C)) and reversed-phase chromatography (RPC) (Figure S5) of purified Ttz-MIC7 revealed a full shift of the antibody peak towards later elution times, with a small peak assigned to an antibody with 1 and a second, main peak to the conjugate with 2 VHH molecules. Liquid chromatography-mass spectrometry (LC-MS) confirmed efficient conjugation of 1 VHH to each antibody HC but no conjugation to the light chain (LC).

**Table 1.**
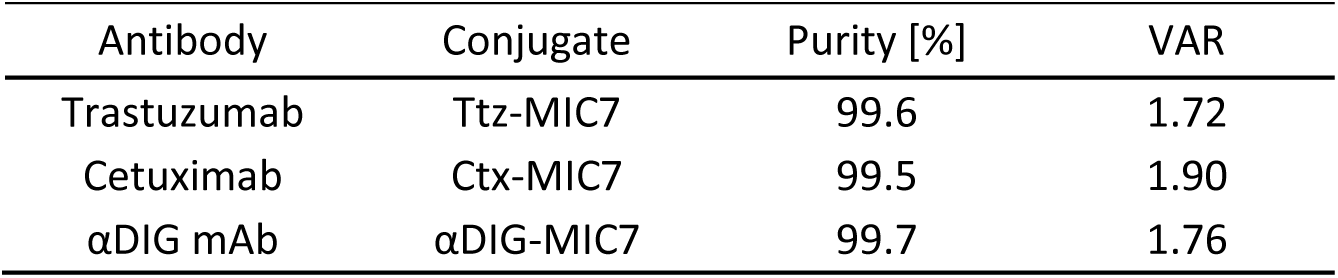
Key data of Ttz-MIC7, Ctx-MIC7 and αDIG-MIC7 antibody conjugates. VHH-to-antibody ratios (VAR) were determined using LC-MS and purity values using analytical SEC.

The LC-MS peak intensities were used to quantify the VHH-to-antibody ratios (VAR) resulting in 1.72 for Ttz-MIC7, 1.90 for Ctx-MIC7 and 1.76 αDIG-MIC7 (Table 1 and Figure S7) which confirms high conjugation efficiencies close the theoretical maximum of 2.

Next, we wanted to evaluate whether our approach is flexible in the choice of antibody and therefore applicable to generate PROTAC shuttles from other IgG1 antibodies. Therefore, we selected additional commercially available antibodies to attach the VHH G_3_-MIC7 and included the antibodies used before for head-to-head comparison. SDS-PAGE analysis of reaction mixes (Figure S9) revealed efficient VHH attachment to the HC of all tested antibodies Ttz, Ctx, Matuzumab, Atezolizumab, Avelumab, Pertuzumab, Rituximab, and αDIG, with VAR values determined by LC-MS ranging from 1.62 to 1.97 (Table 2). SEC analysis showed a high monomer content for all reaction mixes indicating no aggregation issues.

**Table 2.** Generation of PROxAb shuttles from several native mAbs and confirmation of conjugation site. G_3_-MIC7 was conjugated using MTG and reaction mixes analyzed by LC-MS and SEC revealing efficient conjugation with VHH-to-antibody ratios (VAR) close to 2 and high purity. Trastuzumab Q295 with a glutamine to alanine substitution at position 295 was not conjugated confirming this position as the conjugation site.

| Antibody | VAR | Purity [%] |
| --- | --- | --- |
| Trastuzumab | 1.88 | 97.8 |
| Cetuximab | 1.97 | 97.2 |
| Matuzumab | 1.91 | 96.7 |
| Atezolizumab | 1.62 | 98.0 |
| Avelumab | 1.96 | 97.4 |
| Pertuzumab | 1.93 | 97.4 |
| Rituximab | 1.95 | 96.4 |
| αDIG IgG1 | 1.84 | 97.4 |
| Trastuzumab Q295A | 0 | 98.5 |

These results demonstrate that the presented approach can serve as a versatile technology for site-specific protein-protein conjugation and the efficient generation of antibody-VHH conjugates. While we demonstrate conjugation of the PROTAC-binding VHH only, this approach is not limited to the generation of PROxAb shuttle conjugates but can probably also serve as a modular platform to produce bispecific antibodies in general. Combination of VHHs of different target specificities with antibodies of additional specificities would allow to quickly generate combinatorial libraries of bispecific antibodies to screen for optimal target combinations. The need for scalable and robust methods to screen bispecific antibody constructs was recently addressed in the “AJICAP second generation” strategy, which employs affinity peptides pre-orientating amine reactive groups to attach Fab fragments to antibodies.^38^ However, this is a two-step synthesis which requires to eliminate excess reaction compounds in between. Alternative approaches for generating dual binding constructs include intein-based conjugation^39^, the SypTag/SypCatcher approach^40^, SynAbs strategy^41^ or controlled Fab arm exchange^42^. Yet, in contrast to our approach, these further technologies are limited to the use of antibodies or Fc fragments equipped with special functionalities and cannot be applied to off-the-shelf antibodies.

### Antibody VHH bioconjugation does not impact cellular binding and uptake of antibodies as well as VHH-PROTAC binding

To evaluate if the antibody binding properties are impaired by the VHH attachment, binding kinetics of Ttz and Ctx as well as respective MIC7-conjugates were measured using biolayer interferometry (BLI). Very similar dissociation constants (Table S1) and binding curves (Figure S10) were recorded for the individual antibodies and corresponding conjugates. Additionally, binding of Ttz and Ttz-MIC7 to target expressing cells was assessed using two HER2-positive (HER2_pos_) cell lines, namely SKBR3 and BT474^43^, and one HER2 low (HER2_low_) expressing cell line, MDA-MB-435^44^. Cells were incubated with Ttz-MIC7, Ttz or isotype control αDIG-MIC7, stained with an AF488-labeled anti-human IgG1 detection antibody and analyzed using flow cytometry.

Much higher mean fluorescence intensities (MFI) were observed for cells incubated with Ttz or Ttz-MIC7 than with control molecules and signal intensities increased with HER2 expression levels reported in literature (SKBR3 > BT474 > MDA-MB-435)^44^ (Figure 3). In addition, no differences between Ttz and the Ttz-MIC7 incubated cells were observed which, together with the BLI kinetic data, demonstrates that the conjugated VHH does not influence the antibody binding properties. Next, we studied the uptake and accumulation of VHH-conjugated antibodies into cancer cells to assess their suitability for intracellular PROTAC delivery. For this, Ttz, Ttz-MIC7 and αDIG-MIC7 were labeled with pH-sensitive pHAb amine reactive dye (Promega) which shows increased fluorescence at acidic pH and thereby allows to detect labeled constructs in acidic cellular compartments such as the lysosome.^45^ The labeling ratios of the conjugates and respective controls ranged between 6.5 and 9.3 dyes per antibody (Table S3) and live cell fluorescence was monitored over a 24-hour period. Relative fluorescent units normalized to cell count and labeling ratio of each construct increased over time in the Ttz and Ttz-MIC7 treated wells (Figure S11 and S12) whereas no fluorescence was observed with the HER2_low_ cell line. Enhanced cellular uptake of Ttz-MIC7 clearly correlates with a higher count of HER2 receptors on SKBR3 compared to a lower HER2 count on BT474 cells.^46^ In summary, efficient and selective cellular uptake of PROxAb shuttles could be demonstrated.

**Figure 3.**
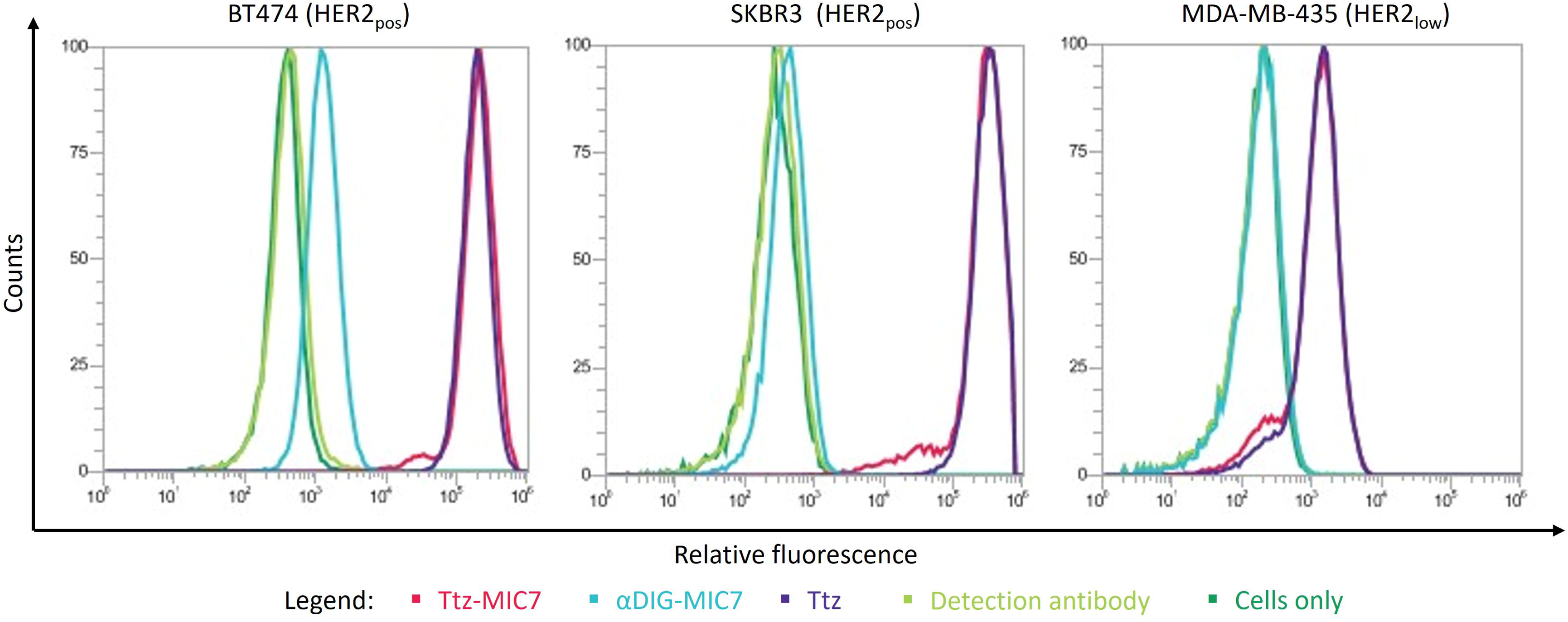
Target cell binding and internalization of conjugated Ttz-MIC7. HER2_pos_ BT474 and SKBR3 cells as well as HER2_low_ MDA-MB-435 cells incubated with Ttz (purple) or Ttz-MIC7 (red) and a fluorescent secondary antibody show equal MFIs compared to controls (isotype control αDIG-MIC7 (cyan), AF488-labeled anti-human detection antibody (light green) and cells only (green)) in flow cytometry analysis.

Finally, we assessed whether conjugation to the antibody altered the binding affinity of MIC7 VHH to PROTACs. Therefore, Ttz-MIC7 and Ctx-MIC7 along with unmodified Ttz and Ctx were immobilized on surface plasmon resonance (SPR) sensors, and their binding kinetic and affinity parameters to the VHL-based PROTACs GNE987, GNE987P and ARV771 were measured. Unmodified antibody controls showed no interaction with PROTACs, while the affinities of Ttz-MIC7 and Ctx-MIC7 to GNE987, GNE987P and ARV771 were in single-digit to sub-nanomolar range (Table S2), which is in line with previous reports for genetically fused PROxAb Shuttles.^23^ These results indicate, that the conjugation of MIC7 via its N-terminus does not affect its affinity to VHL based PROTACs which is in line with literature findings reporting no loss of PROTAC binding after genetic fusion to antibody C-termini.^42, 47, 48^

### Incubation of conjugated PROxAb shuttles with PROTACs results in formation of defined complexes

Next, we assembled PROxAb shuttles via incubation of VHH-antibody conjugates with BRD4 degrading PROTACs. Therefore, Ttz-MIC7 was incubated with increasing concentrations of GNE987, and complex formation was monitored by HIC and expressed as PROTAC to antibody ratio (PAR). Addition of 0.5 molar equivalents of PROTAC per conjugated VHH resulted in the appearance of two new peaks at later elution times interpreted as complexes with 1 (P1) or 2 PROTACs (P2) per antibody (Figure 4). Increasing equivalents (1:1 to 1:2) of added PROTAC resulted in a stepwise decrease of the uncomplexed species (P0) accompanied by an increase in P2 species but no appearance of higher loaded species indicating successful formation of defined PROxAb shuttle complexes carrying 1 PROTAC per VHH.

**Figure 4.**
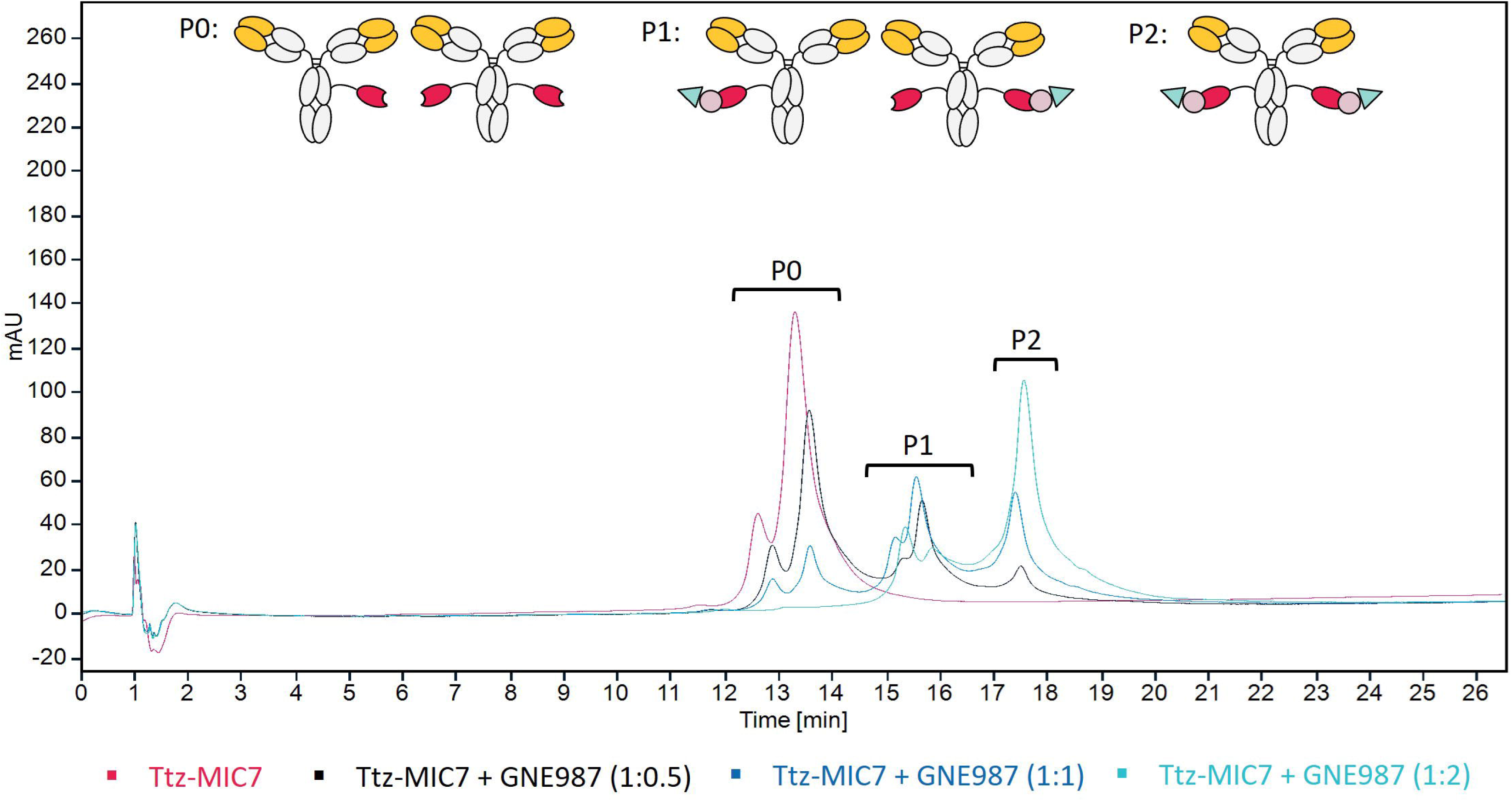
Assembly of PROxAb shuttle-PROTAC complexes. Incubation of Ttz-MIC7 with GNE987 at molar ratios of 1:0.5, 1:1 to 1:2 followed by monitoring of complex formation via HIC. P0, P1 and P2 indicate the elution areas corresponding to PROxAb shuttle conjugates loaded with 0, 1, or 2 PROTACs, respectively.

Following these initial experiments, complex formation of Ttz-MIC7, Ctx-MIC7 and αDIG-MIC7 in combination with GNE987 and GNE987P was investigated in a similar fashion. HIC peak areas were used to determine the PAR. To judge on the efficiency of PROTAC complexation, the VARs (which equal the theoretical maximum PAR assuming each VHH can bind one PROTAC molecule) were divided by the respective PAR resulting in the loading efficiency (in detail see supplemental section 6). As summarized in Table 3, all conjugates incubated with PROTACs at a 1:2 ratio efficiently formed anticipated complexes with loading efficiencies ranging from 73 % to near complete loading resulting in PARs ranging from 1.39 to 1.73.

**Table 3.**
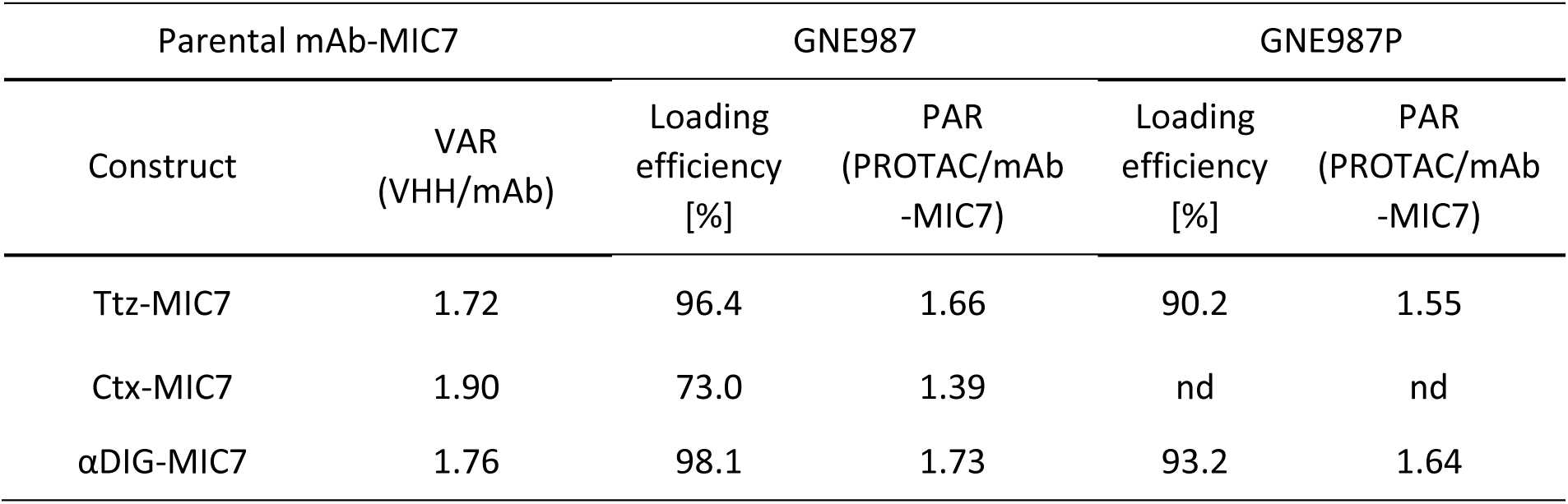
Efficiency of PROTAC complexation. PAR determined by HIC after incubation of antibody-VHH conjugates and PROTACs at a 1:2 molar ratio. Loading efficiencies were calculated by dividing PAR and VAR values. nd: not determined.

| Parental mAb-MIC7 |  | GNE987 |  | GNE987P |  |
| --- | --- | --- | --- | --- | --- |
| Construct | VAR<br>(VHH/mAb) | Loading<br>efficiency<br>[%] | PAR<br>(PROTAC/mAb<br>-MIC7) | Loading<br>efficiency<br>[%] | PAR<br>(PROTAC/mAb<br>-MIC7) |
| Ttz-MIC7 | 1.72 | 96.4 | 1.66 | 90.2 | 1.55 |
| Ctx-MIC7 | 1.90 | 73.0 | 1.39 | nd | nd |
| αDIG-MIC7 | 1.76 | 98.1 | 1.73 | 93.2 | 1.64 |

### PROxAb shuttles complexed with PROTACs mediate cell specific BRD4 degradation and cell killing

Finally, *in vitro* cell based assays were performed to evaluate the biological activity of BRD4 degrading PROTACs complexed with PROxAb shuttles. Degradation was evaluated in SKBR3 cells treated for 24 hours with GNE987P-loaded conjugates, free GNE987P or Ttz-MIC7 alone. Following cellular fixation and permeabilization, the BRD4 level was assessed using an anti-BRD4 antibody and fluorescence-labeled detection antibody by immunofluorescence assay. Thus, the fluorescence intensity (green signal) correlates with the BRD4 level.

Complexed GNE987P and free GNE987P effectively induced BRD4 degradation at nanomolar concentrations as judged from a concentration-dependent reduction in green signal. In contrast, unloaded Ttz-MIC7 and non-binding control αDIG-MIC7 loaded with GNE987P had no impact on BRD4 levels (Figure 5 (A)), indicating that degradation is indeed triggered by selectively delivered PROTAC. In contrast, no degradation effect observed for αDIG-MIC7 + GNE987P indicating that GNE987P is efficiently bound to αDIG-MIC7 which prohibits it from entering the cells. Similarly, effective BRD4 degradation in the nanomolar range was obtained upon treatment with GNE987 and Ttz-MIC7 + GNE987 (Figure S13). In contrast to treatment with free PROTAC, where BRD4 degradation was observed in the double-digit nanomolar range, Ttz-MIC7 complexed PROTACs induced reduced BRD4 levels already at single-digit concentrations. In line with our data, Maneiro and colleagues observed increased BRD4 degradation in immunofluorescence assays after targeted delivery of GNE987.^18^

**Figure 5.**
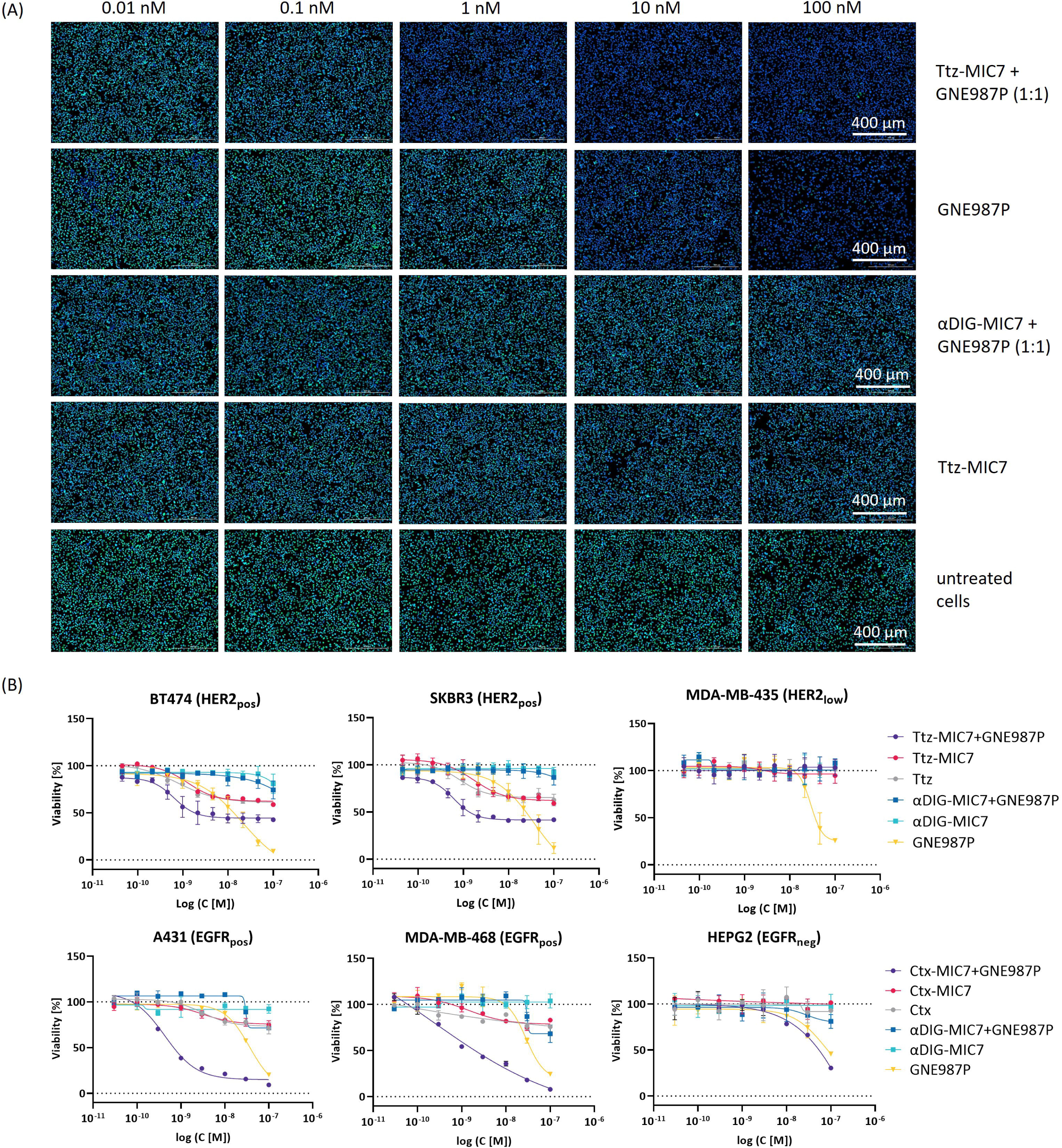
Targeted BRD4 degradation and in vitro potency. (A) Monitoring of BRD4 degradation level in SKBR3 (HER2_pos_) cancer cells via immunofluorescence microscopy analysis. Cells were treated with Ttz-MIC7 or aDig-MIC7 complexed with GNE987P (molar ratio 1:1) or controls for 24 h and afterwards fixated, permeabilized and stained with primary anti-BRD4 antibody, secondary Alexa Fluor™ 488-labeled antibody (green.) Cell nuclei are counterstained with Hoechst dye (blue). (B) Dose-response curves of target positive and low/negative cell lines treated with Ttz-MIC7, Ctx-MIC7 and aDig-MIC7 complexed with GNE987P (molar ratio 1:1) and control molecules. Cells were treated for six (upper panel) or three days (lower panel). Each curve represents the mean of technical duplicates, standard deviation shown as error bars.

As treatment with BRD4-degraders is known to affect cellular viability, we assessed our constructs in *in vitro* cell potency assays. HER2_pos_ (BT474, SKBR3) and HER2_low_ (MDA-MB-435) cells were treated with serial dilutions of Ttz-MIC7 + GNE987P, αDIG-MIC7 + GNE987P and free GNE987P for six days followed by determination of cell viability using CellTiter Glo® (Promega) luminescence assay. Free PROTAC GNE987P reduced viability of all cells irrespective of HER2 expression by 80-90 % with half maximal inhibitory concentration (IC_50_) values in the single digit nanomolar range. In contrast, Ttz-MIC7 + GNE987P showed sub-nanomolar potency on HER2_pos_ cell lines with viabilities reduced by about 60 % while no effects were seen on HER2_low_ cells (Figure 5 (B), Table S4) suggesting increased and specific potency on target positive cells only. Additionally, Ctx-based molecules as well as controls were tested on EGFR_pos_ (A431, MDA-MB-468) and EGFR_low_ (HEPG2) cancer cells. Cell viabilities determined after three days treatment revealed selective cell killing of both EGFR_pos_ cell lines (A431, MDA-MB-468) at sub-nanomolar concentrations and no cytotoxic effect within the evaluated concentration range on EGFR_low_ (HEPG2) cells (Figure 5 (B) and Table S4). Notably, non-binding control αDIG-MIC7 mixed with GNE987P did not induce significant reduction of cell viability in all assays confirming that PROxAb complexing strongly reduces the PROTAC activity on cells that do not express the antibody target. Consistent with previous reports, we observed anti-proliferative effects for Ttz and Ctx antibodies as well as unloaded conjugates.^49, 50^ Equivalent experiments using PROTAC GNE987 were conducted as well and also revealed selective killing of target cells, however free GNE987 was more potent than GNE987P and complexed shuttles showed similar potencies on target cells as GNE987 (Table S4). In contrast, potency of complexed GNE987P was increased on target-positive cells by about ten- to 100-fold compared to PROTAC alone, which is likely due to the lower cellular permeability of GNE987P that is overcome by the shuttle-mediated transport.^34^ Similar effects on target cells where observed previously by Pillow et al who covalently conjugated GNE987 to an anti-CLL1 antibody.^33^

In conclusion, the tested mAb-VHH conjugates show very similar characteristics as the original PROxAb shuttle molecules allowing for complexation of PROTACs and specific delivery to target cells only.^23^ However, while the original technology requires individual recombinant production of each mAb-VHH fusion protein, the technology presented herein can be used for straight-forward generation of PROxAb shuttles from off-the-shelf antibodies to rapidly screen for the most promising mAb-VHH combinations.

## Conclusion

In summary, we have developed a new approach for effective chemoenzymatic conjugation of VHH to native antibodies. Through fusion of an N-terminal recognition tag to VHH we were able to site-specifically conjugate the VHH to glutamine 295 of the heavy chain in native IgG-based antibodies and thereby generate bispecific antibodies. This novel approach was exemplarily used to generate PROxAb Shuttle conjugates by coupling G_3_-MIC7 VHH with native IgG-based antibodies. The biological functionality of the conjugate was demonstrated by loading with BRD4 degrading PROTAC and subsequent immunofluorescence degradation and cell viability experiments. This chemoenzymatic method is a modular approach that could be applied for high-throughput screening to identify ideal binder combinations and fits into both worlds, the fast growing field of TPD and bispecific antibodies.

## Notes

H.S., J.T., J.S., D.S., C.S., S.H., J.A., N.R., M.R. and S.D. are employees of Merck KGaA, Darmstadt, Germany.

## Acknowledgements

The authors kindly like to thank L. Basset, M. Fleischer, J. Finkernagel, J. Hannewald, S. Jäger, S. Keller, V. Lautenbach, K. Leidinger, I. Schmidt, A. Schoenemann, P. Steiner, and K. Waurisch for advice and laboratory support.

## Abbreviations

ADC: antibody-drug conjugates
TPD: targeted protein degradation
PROTACs: proteolysis-targeting chimeras
UPS: ubiquitin-proteasome system
CRBN: Cereblon
VHH: camelid single-domain antibody
VHL: Von Hippel-Lindau
MTG: microbial transglutaminase
BRD4: bromodomain-containing protein 4
HER2: human epidermal growth factor 2
Ttz: Trastuzumab
Ctx: Cetuximab
HIC: Hydrophobic Interaction Chromatography
RPC: Reversed-Phase Chromatography
VAR: VHH-to-antibody ratio
LC-MS: liquid chromatography-mass spectrometry
αDIG: digoxigenin targeting human IgG1 antibody
MFI: Mean fluorescence intensity
RFU: relative fluorescent unit
mAb: monoclonal antibody
HC: heavy chain
SPR: Surface Plasmon Resonance

