## Supporting Information for "Welding PROxAb Shuttles: A modular approach for generating bispecific antibodies via site-specific protein-protein conjugation"

[‡] Merck KGaA, Frankfurter Str. 250, 64293 Darmstadt, Germany

Table of Contents

Experimental procedures ........ - 2 -

1. Amino acid sequence of $G_3$-MIC7 for MTG mediated conjugation ........ - 2 -
2. Recombinant expression and purification of $G_3$-VHH ........ - 2 -
3. Protein-protein conjugation ........ - 2 -
4. Assembly of PROxAb shuttle-PROTAC complexes ........ - 2 -
5. Analytical SE-HPLC ........ - 3 -
6. RP-HPLC analysis ........ - 3 -
7. Determination of shuttle-PROTAC complexation using HIC ........ - 3 -
8. VAR determination using LC-MS ........ - 3 -
9. Bio-layer-Interferometry for characterization of antigen-binding of the antibody domains- 4 -
10. Surface plasmon resonance spectroscopy for characterization of PROTAC-binding of the VHH antibody domains ........ - 4 -
11. Cell culture ........ - 4 -
12. Fluorescence activated cell sorting analysis ........ - 4 -
13. Live-cell imaging of conjugated PROxAb shuttle internalization ........ - 5 -
14. Immunofluorescence assay to determine BRD4 degradation ........ - 5 -
15. Cytotoxicity assay ........ - 5 -

Supplementary Figures and Tables ........ - 6 -

References ........ - 14 -

Experimental procedures

1. Amino acid sequence of $G_3$-MIC7 for MTG mediated conjugation

| | | | | | |
|---|---|---|---|---|---|
| 1 | *RGRKLQDRLA* | *KEGMHSSALL* | *CCLVLLTGVR* | *A* **GGG** GSGGGS | GGSGGGGSGA |
| 51 | VQLVESGGGL | VQAGGSLRLS | CAASGFSFDD | YALGWFRQAP | GKEPEGLSCI |
| 101 | SSSDGSTWYA | DSVKGRFTIS | SDNAKNTVYL | QMNSLKPEDT | AVYYCSAIYR |
| 151 | LSCSVVRPTI | RYALDYWGQG | TQVTVSS | | |

*Signal peptide,* **MTG recognition tag**, glycine-serine spacer, $G_3$-MIC7 VHH [1]

2. Recombinant expression and purification of $G_3$-VHH

A plasmid based on pcDNA 3.4 for recombinant expression of VHH $G_3$-MIC7 was generated by Biointron, China, and used for transfection of Expi293™ cells using the Expression System Kit (Thermo Fisher) according to the manufacturer's protocol. Transfected cells were harvested 6 days post transfection via centrifugation at 3000 rpm for 20 minutes at 4 °C. Supernatant was filtered through 0.2 µm PES filter (Nalgene) before purification. The VHH was purified by Amsphere™ A3 affinity chromatography (JSR Life Sciences) followed by buffer exchange with HiPrep™ 26/10 (Cytiva), concentrated using Ultra-15 centrifugal filter devices (molecular weight cutoff (MWCO) 3000, Amicon) and sterile filtered (0.2 µm). Protein concentrations were determined by NanoDrop One UV-Vis Spectrophotometer (Thermo Fisher) at 280 nm using the extinction coefficient calculated from the amino acid sequence. The VHH purity was analyzed using analytical SE-HPLC and SDS-PAGE gel electrophoresis (Invitogen); protein identity as well as full cleavage of signal peptide were confirmed by intact mass analysis by LC-MS using TripleTOF 6600+ mass spectrometer (AB Sciex).

3. Protein-protein conjugation

Microbial transglutaminase (MTG) was produced as described before[2] and the enzyme activity was determined via photometric assay ZediXclusive (Zedira). Native antibodies were either used as clinical grade material (trastuzumab, cetuximab, avelumab, pertuzumab, rituximab, matuzumab) or produced as described elsewhere (αDig, atezolizumab, trastuzumab Q295A).[3] The MTG-mediated antibody conjugation was performed with 5 mg/mL antibody, 10 molar equivalent VHH and 10 U/mL MTG. Reaction mixes were incubated for 24 hours at 30 °C and 550 rpm, stopped with MTG blocker C102 (Zedira) or directly purified by preparative size exclusion chromatography (SEC). Purification was performed in a HPLC 1260 system (Agilent Technologies) using Superdex 200 10/300 GL increase column and running buffer PBS pH 7.0. Fractions of expected elution time were pooled, concentrated using Ultra-15 centrifugal filter devices (MWCO 50000, Amicon) and sterile filtered (0.2 µm). Protein concentrations were measured by NanoDrop One UV-Vis Spectrophotometer (Thermo Fisher) at 280 nm and determined using the theoretical extinction coefficient calculated based on the amino acid sequence. Purified samples or stopped reaction mixes were analyzed by SE-HPLC and SDS-PAGE (Invitrogen), and VHH-to-antibody ratio (VAR) was determined by HIC, RP-HPLC and LC-MS.

4. Assembly of PROxAb shuttle-PROTAC complexes

PROxAb shuttles-PROTAC complexes were assembled by incubating 10 µM VHH-antibody conjugates with differing molar equivalents of VHL-based PROTACs (GNE987 (MedChemExpress), GNE987P (MedChemExpress) or ARV771 (MedChemExpress)) in PBS pH 7.4 with 5 % DMSO at 25 °C, 650 rpm for 3 hours. For cellular experiments, 0.3 % Tween-20 (Merck KGaA) final were added to enable sample application to cells by Tecan D300e dispenser (Tecan Group Ltd.).

5. Analytical SE-HPLC

Analytical size exclusion-HPLC (SE-HPLC) was performed using a Bio Resolve SEC mAb column (4.6 x 150 mm, 2.5 µm, 200 Å, Waters) with 0.05 M sodium phosphate, 0.4 M sodium perchlorate, pH 6.3 (Merck KGaA) as running buffer. A flow rate of 0.35 mL/min and a total run time of 7 min were used. Typically, 10 µg sample were injected, and absorption signal recorded at 214 nm.

6. RP-HPLC analysis

Reversed Phase-HPLC (RP-HPLC) was performed with a PLRPS column (2.1 x 50 mm, 5 µm, 4000 Å, Agilent technologies) at 65 °C. A gradient of 30 to 45 % solvent B was run over 7.5 min at a flow rate of 1 mL/min. 10 µg sample were injected, and absorption signal recorded at 214 nm. The solvents are as follows: Solvent A: water containing 0.1 % (v/v) TFA , solvent B: acetonitrile containing 0.1 % (v/v) TFA (Merck KGaA).

7. Determination of shuttle-PROTAC complexation using HIC

Hydrophobic interaction chromatography (HIC) analysis was performed with a TSKgel® Butyl-NPR column (4.6 x 100 mm, 2.5 µm, Tosoh bioscience) heated to 35 °C. Samples were adjusted to 0.5 M ammonium sulphate and 42.5 µg sample were injected per run. A gradient ranging from 40 % to 80 % over 40 minutes at a flow rate of 0.45 mL/min was applied, starting from 0.025 M Tris HCl (Merck KGaA) pH 7.5 to a mixture of 20 % isopropanol (Merck KGaA), 0.025 M Tris HCl, 2 M ammonium sulfate (Merck KGaA) pH 7.5. Previously, Absorption was recorded at a wavelength of 280 nm.

The efficiency of PROTAC loading was calculated as follows:

*Equation 1*

$$Loading\ efficiency = \frac{Area_{P0} * 0\ +\ Area_{P1} * 1\ +\ Area_{P2} * 2}{VAR * Area_{total}}$$

While $Area_{P0}$ is the area of the peak referred to PROxAb Shuttle without any bound PROTAC, $Area_{P1}$ the area of the peak referred to PROxAb shuttle with one PROTAC molecule and $Area_{P2}$ to the area of the peak referred to PROxAb shuttle with two PROTAC molecules. VAR is the VHH-to-antibody ratio, and $area_{total}$ the total area of all peaks in the chromatogram.

8. VAR determination using LC-MS

For determination of the VHH-to-antibody ratio (VAR) using LC-MS, conjugates were diluted with 0.1 % formic acid (Merck KGaA) to a final concentration of 0.05 mg/ml. Subsequently, 100 µL of this solution were reduced with 1 µL TCEP (500 mM) for 5 min at room temperature. LC-MS analysis was performed using an Exion HPLC system (buffer A: 0.1 % formic acid in water, buffer B: 0.1 % formic acid in acetonitrile) coupled to a Sciex 6600+ mass spectrometer by a Turbo V ESI source. 4 µl protein solution was loaded onto a bioZEN 3.6 µm Intact C4, 2.1 x 50 mm column (Phenomenex) and eluted with a linear gradient from 15 % to 95 % buffer B within 3 min at 0.25 min/ml flow rate. Data were acquired with positive polarity and in a mass range from 400 to 4000 m/z. Other instrument settings were as follows: source voltage 5.5 kV, declustering potential 180 V, accumulation time 1 s, source temperature 350 °C, gas1 50 l/h, gas2 25 l/h, and curtain gas 10 l/h. The mass spectrometer was calibrated with ESI positive calibration solution for the SCIEX X500B. Acquired data were processed with Genedata Expressionist 16.5. Spectra were smoothed with the moving average algorithm and baseline subtracted with the quantile method. Spectra were deconvoluted with the maximum entropy method (20 iterations, high quality). For protein mapping the tolerance was set to 10 Da and glutamine to pyroglutamate conversion as well as C-terminal lysine loss were selected as variable modifications.

9. Bio-layer-Interferometry for characterization of antigen-binding of the antibody domains

Bio-layer-Interferometry was performed on Octet RED96 system (ForteBio, Pall Life Science) at 25 °C. Parental antibodies and mAb-VHH conjugates concentrated at 12.5 µg/ml in PBS were loaded on anti-human IgG Fc capture (AHC) sensors (ForteBio) for 180 s, then rinsed in kinetic buffer (KB, PBS + 0.1 % Tween-20 + 1 % bovine serum albumin (BSA, Merck KGaA)) for 45 s and associated with the corresponding antigen at concentrations ranging from 15.6 nM to 500 nM for 300 s. Subsequent dissociation was performed in KB for 600 s. The association in pure KB was used as reference value. Data were fitted using ForteBio data analysis software 8.0, the Savitzky-Golay filter and a 1:1 binding model.

10. Surface plasmon resonance spectroscopy for characterization of PROTAC-binding of the VHH antibody domains

The kinetic and affinity parameters of the Ttz-MIC7/PROTAC and Ctx-MIC7/PROTAC interactions were evaluated by surface plasmon resonance (SPR). Ctx, Ctx-MIC7, Ttz or Ttz-MIC7 were immobilized onto a high-capacity amine sensor chip (Bruker Daltonics) via the standard amine coupling procedure at 25 °C. Prior to immobilization, the carboxymethylated surface of the chip was activated with 200 mM 1-ethyl-3-(3-dimethylaminopropyl)-carbodiimide and 25 mM N-hydroxy succinimide (NHS) for 10 min. The Ctx-MIC7 variants were diluted to 10 µg/mL in 10 mM BisTris at pH 6.0 and immobilized on the activated surface chip for 3 to 7 min, in order to reach 3,000 to 9,000 response units (RU). The remaining activated carboxymethylated groups were blocked with 1 M ethanolamine pH 8.0 in a 7 min injection step. HBS-N, which consists of 10 mM HEPES pH 7.4 and 150 mM NaCl, was used as the background buffer during immobilization. PROTACs were prediluted in DMSO (Honeywell), diluted 1:50 in running buffer (12 mM phosphate, pH 7.4, 137 mM NaCl, 2.7 mM KCl, 0.05 % Tween-20, 2 % DMSO) and injected at 10 different concentrations using two-fold dilution series, from 0.5 µM to 0.001 µM. A DMSO solvent correction (1 % - 3 %) was performed to account for variations in bulk signal and to achieve high-quality data. Interaction analysis cycles consisted of a 300 s sample injection (30 µL/min; association phase) followed by 1,800 s of buffer flow (dissociation phase). All sensorgrams were processed by first subtracting the binding response recorded from the control surface (reference flow-channel), followed by subtraction of the buffer blank injection from the active flow-channel (target protein immobilized). All datasets were fit to a simple 1:1 Langmuir interaction model to determine the kinetic rate constants. The experiments were performed on a SPR-32 PRO (Bruker Daltonics, Bremen, Germany) at 25 °C and the interactions were evaluated using the provided Sierra Analyser Software (version 3.4.5.).

11. Cell culture

Human cancer cells were obtained from the American Type Culture Collection (cell lines: SKBR3, BT474, MDA-MB-435, A431, MDA-MB-468, HEPG2) and cultured at 37 °C, 5 % $CO_2$ and 95 % humidity. SKBR3, MDA-MB-468 and HEPG2 cells were cultured in high-glucose Dulbecco's modified Eagle's medium (DMEM, Sigma-Aldrich) containing 10 % fetal bovine serum (FBS, Sigma-Aldrich), A431 cells in Roswell Park Memorial Institute (RPMI, Sigma-Aldrich) 1640 medium with 10 % FBS, 2 mM L-glutamine (Sigma-Aldrich), and 1 mM sodium pyruvate (Sigma-Aldrich), BT474 cells in DMEM/F-12 (Sigma-Aldrich) medium supplemented with 2 nM L-glutamine, 1 nM sodium pyruvate, 1 µg/mL insulin (Sigma-Aldrich) and 10 % FBS and MDA-MB-435 cells in DMEM/F-12 supplemented with 10 % FBS, 2 nM L-glutamine, 1 % NEAA (Sigma-Aldrich) and 1 nM sodium pyruvate.

12. Fluorescence activated cell sorting analysis

HER2 expressing ($HER2_{pos}$) SKBR3, BT474 and MDA-MB-435 ($HER2_{low}$) or EGFR expressing ($EGFR_{pos}$) A431, MDA-MB-468 and HEPG2 ($EGFR_{low}$) were detached, washed twice with cold PBS + 1 % BSA and seeded in 96-well plates (10000 viable cells/well, Greiner). Cells were incubated in duplicates with 100 nM sample (parental antibody or conjugate) for 1 hour on ice and washed twice with cold PBS + 1 % BSA. 500 nM detection antibody (Alexa Fluor® 488 AffiniPure Fab Fragment Goat Anti-Human IgG, Fcγ fragment specific (Jackson ImmunoResearch, Cat: 109-547-003)) was added and incubated for 1 hour on ice. Following two washing steps with PBS + 1 % BSA on ice, cells were resuspended in 200 µL of 5 nM final SYTOX™ Red (Invitrogen™) diluted in PBS + 1 % BSA. Cells treated only with detection antibody served as control, untreated cells were used for normalization of fluorescence signal. Attune™ NxT Flow Cytometer (Thermo Fisher Scientific) with Attune™ Cytometric Software 3.2.1526.0 was used to analyze the events per well.

13. Live-cell imaging of conjugated PROxAb shuttle internalization

SKBR3, BT474 and MDA-MB-435 cells were diluted in their appropriate growth medium (see chapter 11), seeded (10000 viable cells/well) in a black, µClear bottom 384-well plate (Greiner) and cultivated overnight according to standard conditions (see chapter 11). Samples were labeled with pHAb-dye using Promega's labeling kit according to manufacturer's protocol using traditional in-solution conjugation. Briefly, antibodies were incubated with 20 molar excess of amine reactive pHAb dye for 1 hours while shaking at 500 rpm and 25 °C in the dark. Subsequently, a desalting step was performed using Zeba™ Spin Column to remove excess dye. Concentration of dye labeled conjugate and the antibody-dye ratio were calculated as recommended by the manufacturer. pHAb-dye labeled constructs were supplemented with 0.3 % Tween-20 final, diluted to 3 µM in PBS, and added to the cells using a Tecan D300e digital dispenser. Cells were shaken at 450 rpm for 2 min. Images were captured using Cell Imaging Multimode Reader Cytation 5 (BioTek/Agilent) (10x objective, excitation: 531 nm, emission: 593 nm, LED intensity: 10, integration time: 14 msec, camera gain: 24) every 30 min for 24 hours.

14. Immunofluorescence assay to determine BRD4 degradation

SKBR3 cells were seeded in respective medium in black, µClear bottom 96-well plates (Greiner) (50000 viable cells/well), followed by overnight incubation in a humid chamber at 37 °C and 5 % $CO_2$. Serial dilutions of samples were added to the cells using Tecan D300e dispenser followed by normalization of total medium volume. Final concentrations of 0.003 % Tween-20 and 0.05 % DMSO were adjusted with 0.3 % Tween-20 in PBS pH 7.4 and DMSO stock solution. Cells were incubated overnight at 37 °C and 5 % $CO_2$ followed by media removal, washing with DPBS and fixation for 15 minutes by incubating with 2 % (v/v) formaldehyde solution. Afterwards, cells were washed and then permeabilizated using 0.2 % (v/v) Triton X-100 for 10 minutes. Cells were blocked with 3 % (w/v) BSA in DPBS at RT for 60 minutes, incubated with the respective primary antibody at 4 °C overnight followed by staining with secondary antibody for 120 minutes and Hoechst staining for 90 minutes, both in the dark and at room temperature. Final analysis was performed using Cell Imaging Multimode Reader Cytation 5 (BioTek/Agilent) and image processing using BioTek gen5 software.

15. Cytotoxicity assay

Cells were seeded in the respective medium in white 384-well plates (Corning®) (2000 viable cells/well) followed by overnight incubation in a humid chamber at 37 °C and 5 % $CO_2$. Complexation of PROTAC and PROxAb shuttle conjugates was performed as previously described. Serial dilutions of samples were added to the cells using a Tecan D300e dispenser with normalization to the highest application volume for each well. The final concentrations of 0.003 % Tween-20 and 0.05 % DMSO were adjusted with 0.3 % Tween-20 in PBS pH 7.4 and DMSO stock solution. Subsequently, cells were incubated at 37 °C and 5 % $CO_2$. Analysis using CellTiter Glo ® reagent (Promega) was performed after three or six days as described in the manufacturer's protocol. Luminescence signals were measured using an Envision reader (Perkin Elmer), luminescence values normalized to the luminescence of untreated cells and data analyzed using a slope sigmoidal response fit model (GraphPad Prism, GraphPad Software, Inc.).

### Supplementary Figures and Tables

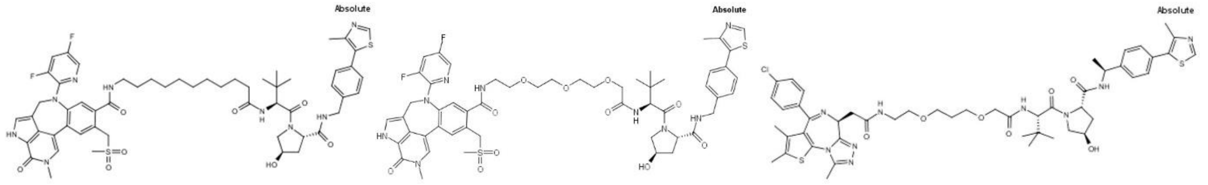


GNE987 GNE987P ARV771

*Figure S 1 Structure of PROTACs GNE987 [4], GNE987P [5] and ARV771 [6].*

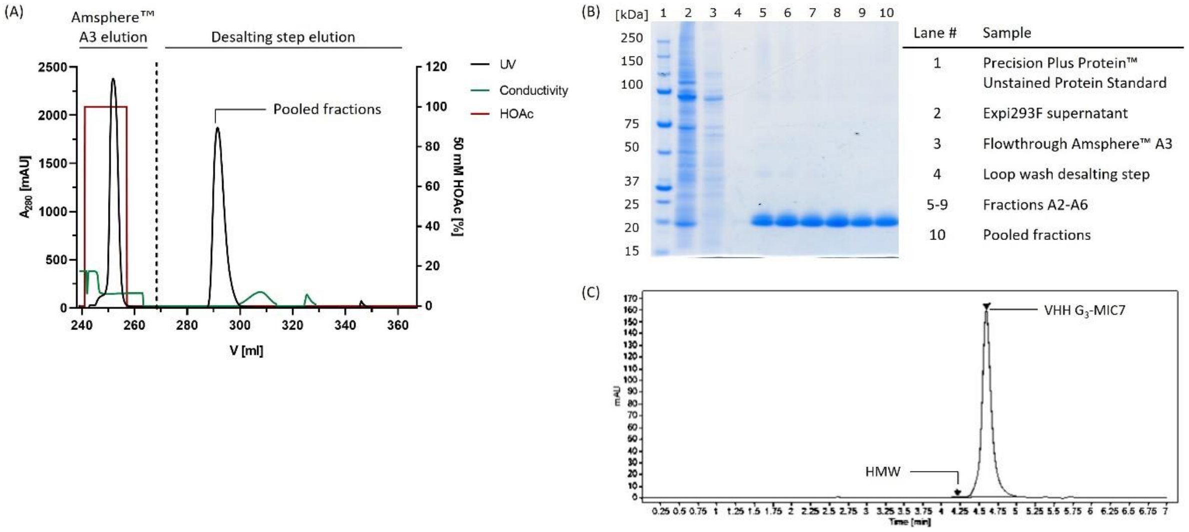


*Figure S 2 Amsphere™ A3 purification process of VHH MIC7. (A) ÄKTA Xpress chromatogram showing the elution of protein in Amsphere™ A3 (1 mL) affinity chromatography with 50 mM acetic acid (HOAc) pH 3.2 and subsequent buffer exchange in PBS pH 7.0 using a HiPrep™ 26/10 desalting column. (B) SDS-PAGE analysis using 4-12 % Bis-Tris gel (Invitrogen) of the following samples: non-reduced Expi293F supernatant, Amsphere™ A3 flowthrough, loop wash of desalting step, fractions A2-A6 and purified VHH MIC7. (C) $A_{280}$ signal of analytical SE-HPLC chromatogram of purified VHH MIC7 shows high purity of 99.3 % and 0.7 % high molecular weight (HMW) species.*

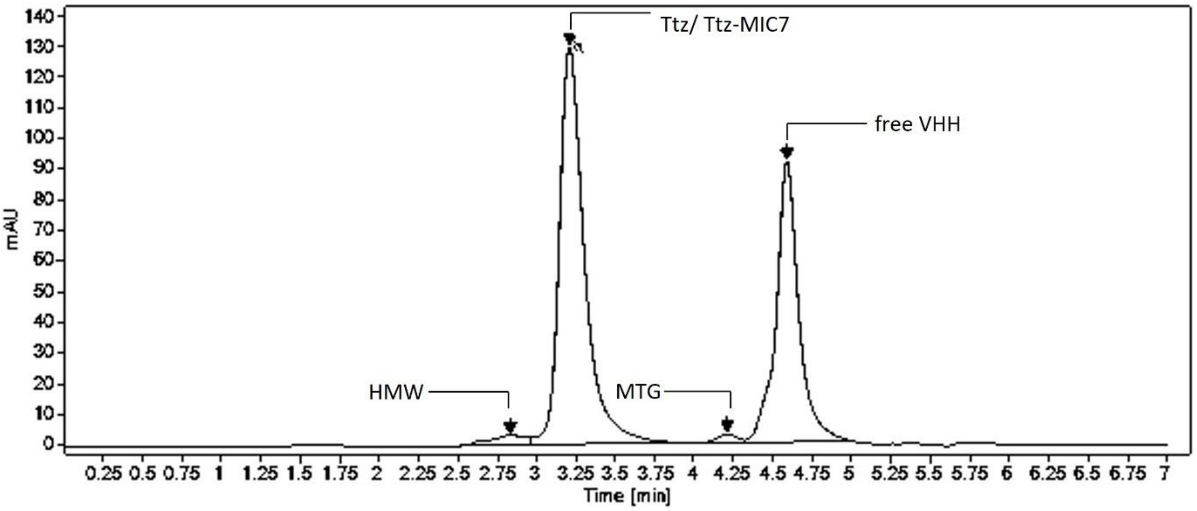


*Figure S 3 $A_{280}$ signal of analytical SE-HPLC chromatogram of conjugated Ttz-MIC7 revealing high purity with low amounts of high molecular weight species (HMW). Unconjugated VHH and MTG elute at 4.25 min and 4.59 min, respectively.*

(A)

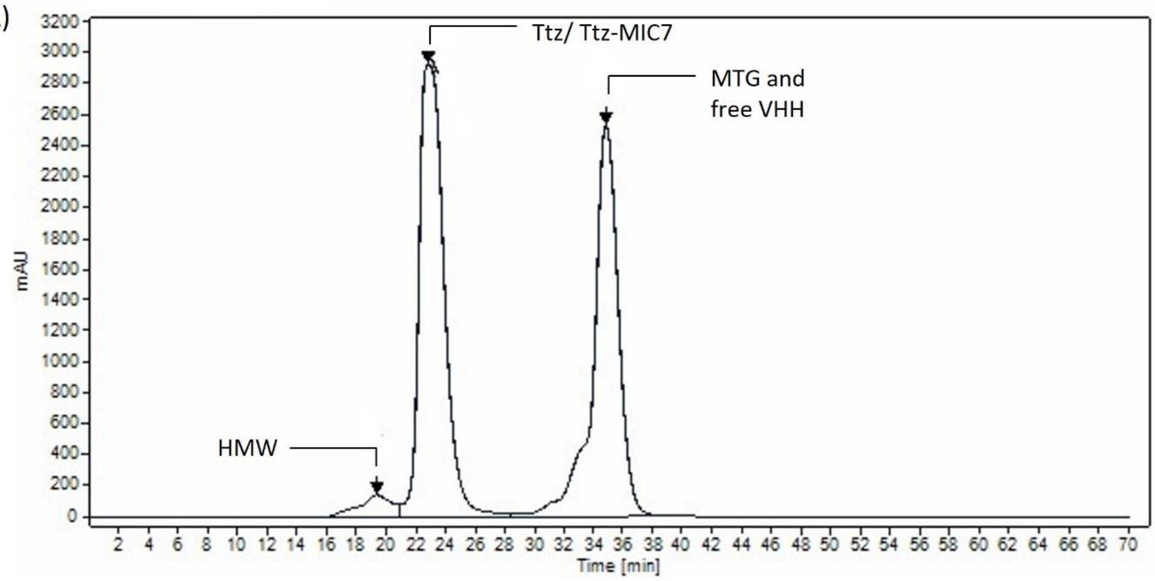


(B)

| Antibody | Yield [mg] | Yield [%] | Purity [%] |
|---|---|---|---|
| Ttz-MIC7 | 1.75 | 87.3 | 99.6 |
| Ctx-MIC7 | 1.46 | 72.8 | 99.5 |
| αDIG-MIC7 | 1.59 | 79.5 | 99.7 |

*Figure S 4 (A) Preparative Size-Exclusion Chromatography (SEC) purification of conjugated PROxAb shuttle, exemplarily shown for Ttz-MIC7. HPLC 1260 system (Superdex 200 10/300 GL increase column, Agilent Technologies) $A_{280}$ chromatogram showing separation of antibody/conjugate peak from HMW, unconjugated VHH and MTG. (B) Final yield and purity (determined by analytical SE-HPLC) after sample concentration and sterile filtration.*

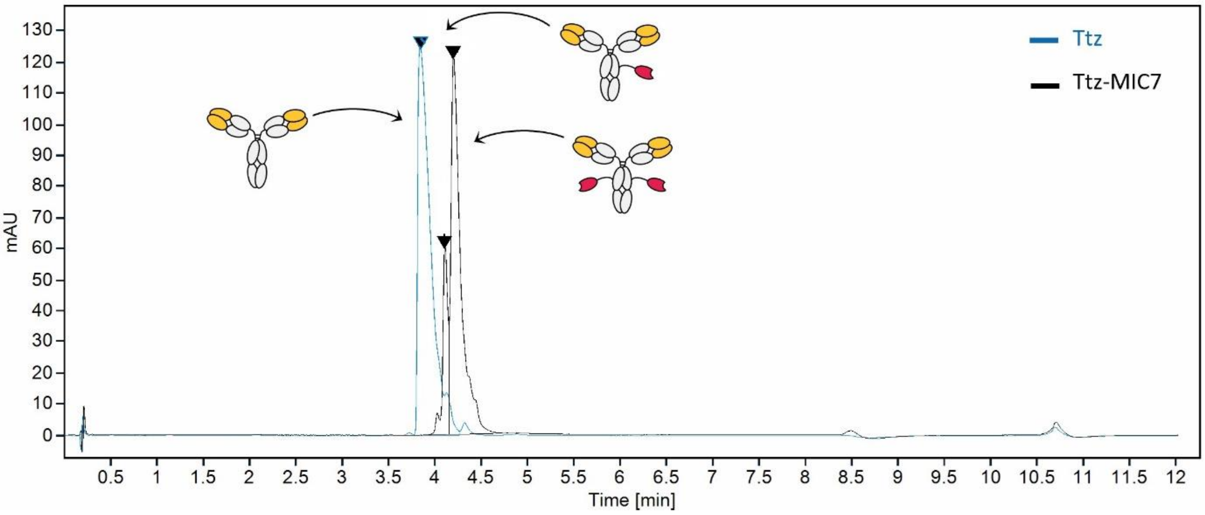


*Figure S 5 Reversed Phase Chromatography (RPC) analysis of mAb-VHH conjugation. Overlay of HPLC 1260 system (TSKgel® Butyl-NPR column, Tosoh bioscience) chromatogram of Ttz (blue) and SEC-purified Ttz-MIC7 (black).*

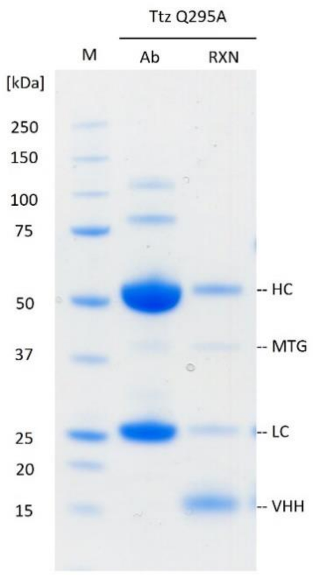


*Figure S 6 Reduced SDS-PAGE of Ttz Q295A conjugation. The parental antibody shows expected bands at 25 kDa (light chain, LC) and 50 kDa (heavy chain, HC). The reaction mix (RNX) shows the same bands plus unconjugated VHH $G_3$-MIC7 (approx. 15 kDa) and MTG (approx.) 38 kDa but no additional bands that would indicate VHH-mAb conjugation.*

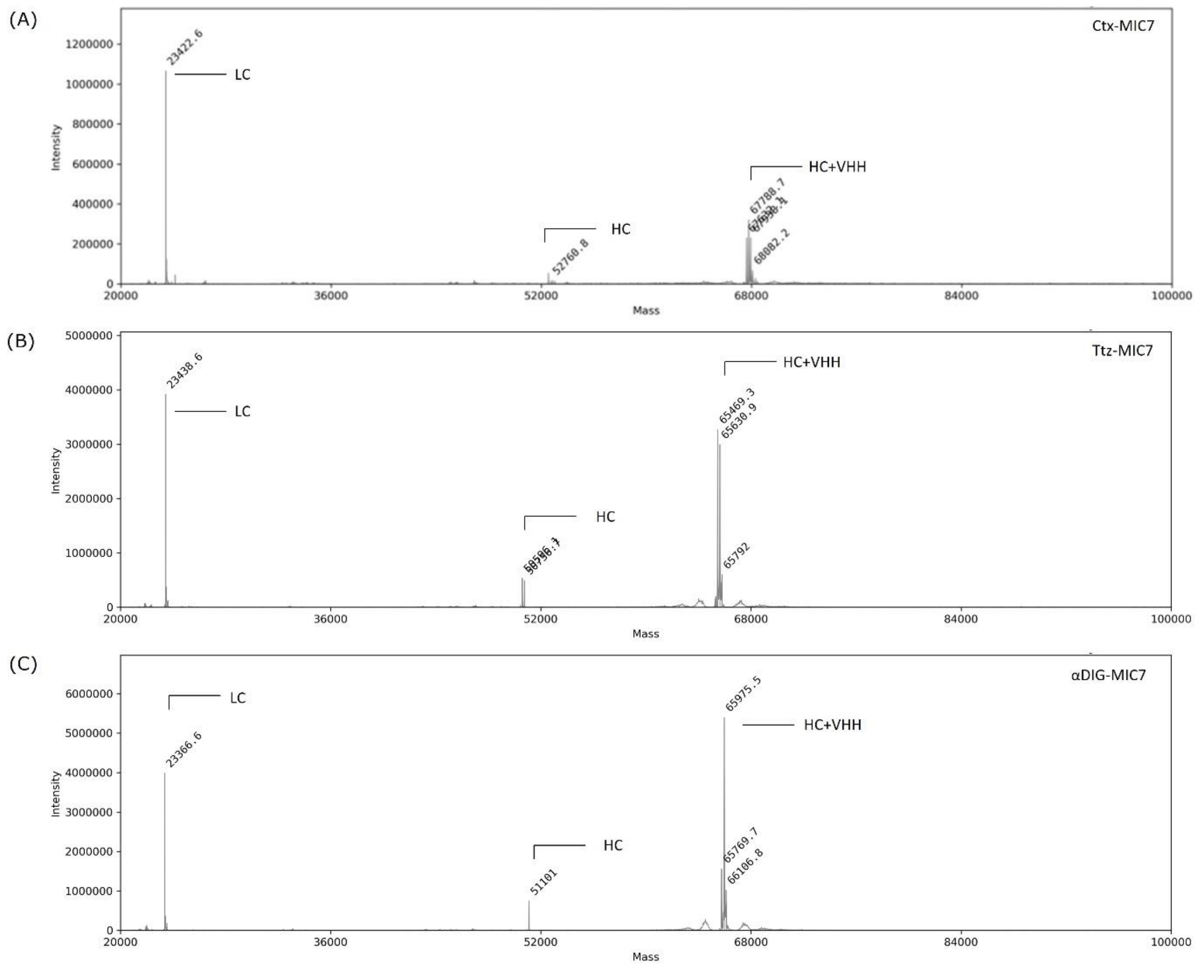


*Figure S 7 VHH-to-antibody ratio (VAR) determination after purification of conjugated PROxAb shuttles. Deconvoluted MS spectra used to assign RP peaks to individual light or heavy chain species conjugated with VHH 'MIC7'. MTG-mediated conjugation to (A) Cetuximab, (B) Trastuzumab or (C) αDIG antibody. Light chain (LC), heavy chain (HC) and HC+VHH were detected.*

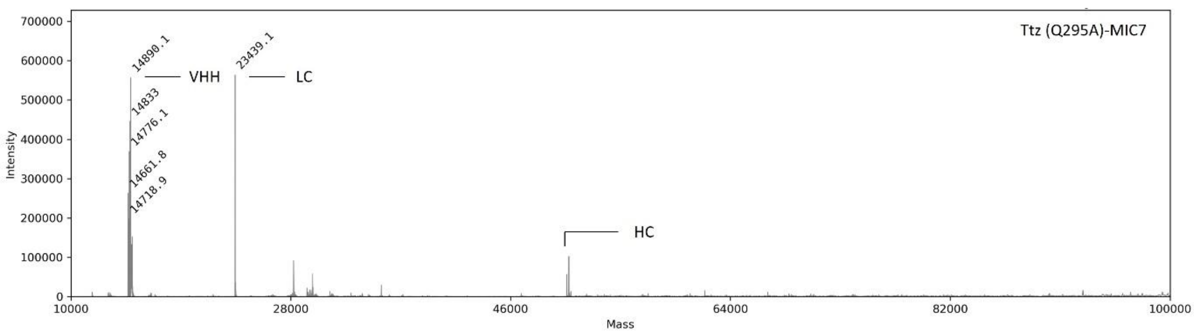


*Figure S 8 VAR determination of conjugated Ttz (Q295A)-MIC7. Deconvoluted MS spectra used to assign PR peaks to individual light or heavy chain species and VHH 'MIC7'.*

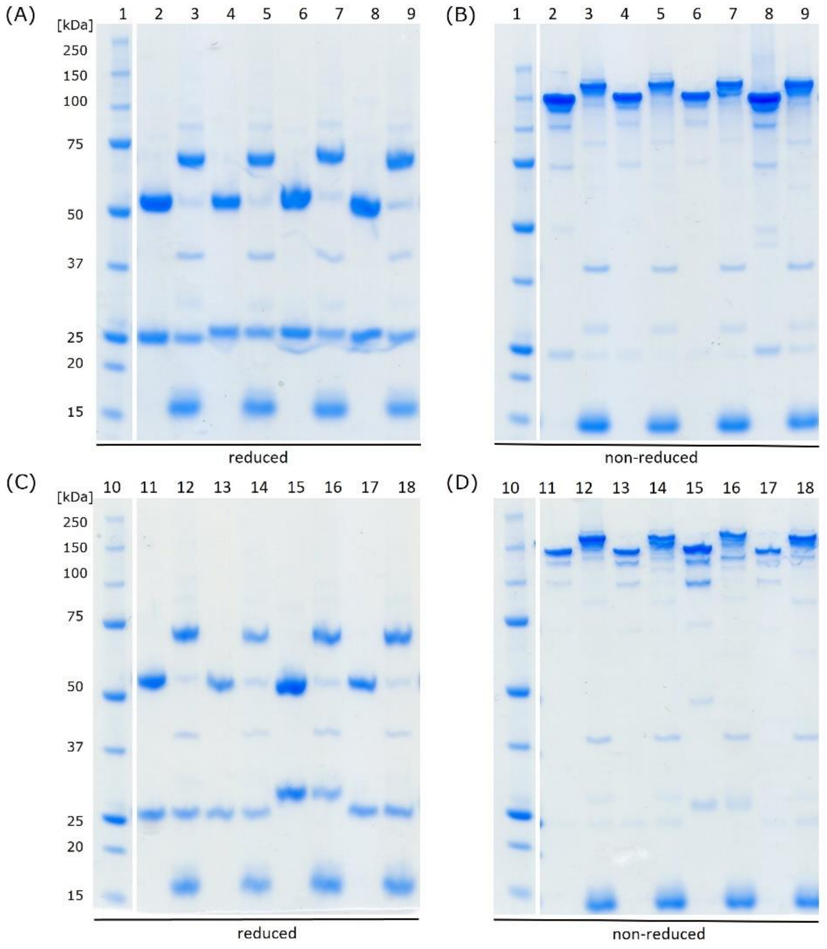


| Lane # | Sample |
|---|---|
| 1 | Precision Plus Protein™ Unstained Standard |
| 2 | Rituximab (Rix) |
| 3 | Conjugate Rix-MIC7 |
| 4 | Ctx PG-LALA |
| 5 | Conjugate Ctx PG-LALA-MIC7 |
| 6 | Cetuximab (Ctx) |
| 7 | Conjugate Ctx-MIC7 |
| 8 | Trastuzumab (Ttz) |
| 9 | Conjugate Ttz-MIC7 |

| Lane # | Sample |
|---|---|
| 10 | Precision Plus Protein™ Unstained Standard |
| 11 | Matuzumab (Mtz) |
| 12 | Conjugate Mtz-MIC7 |
| 13 | Atezolizumab (Atz) |
| 14 | Conjugate Atz-MIC7 |
| 15 | Avelumab (Ave) |
| 16 | Conjugate Ave-MIC7 |
| 17 | Pertuzumab (Ptz) |
| 18 | Conjugate Ptz-MIC7 |

*Figure S 9 SDS-PAGE gels showing reduced (A and C) and non-reduced (B and D) reaction mixes and parental antibodies. Bands in (A) and (C) of parental antibodies appear at expected 25 kDa (LC) and 50 kDa (HC), whereas an expected shift of the heavy chain band toward 65 kDa is visible in the reaction mixtures. Free excess of VHH $G_3$-MIC7 appears at approx. 15 kDa and MTG at 38 kDa, in both non-reduced and reduced reaction mixes. In (B) and (D), the parental antibodies appear at the expected masses of approx. 150 kDa. The bands of the reaction mixtures are distributed among unconjugated 150 kDa (VAR 0) and conjugated 165 kDa (VAR 1) and 180 kDa (VAR 2) species. Ctx PG-LALA refers to cetuximab with the mutation P329G/L234A/L235A. Materials and conditions: 4-12 % Bis-Tris Gel (Invitrogen), MES SDS running buffer (1x), 40 min at 200 V, stained with Der Blaue Jonas (Coomassie-based, German Research Products) for 2 h, marker: Precision Plus Protein™ Unstained Standard (BioRad).*

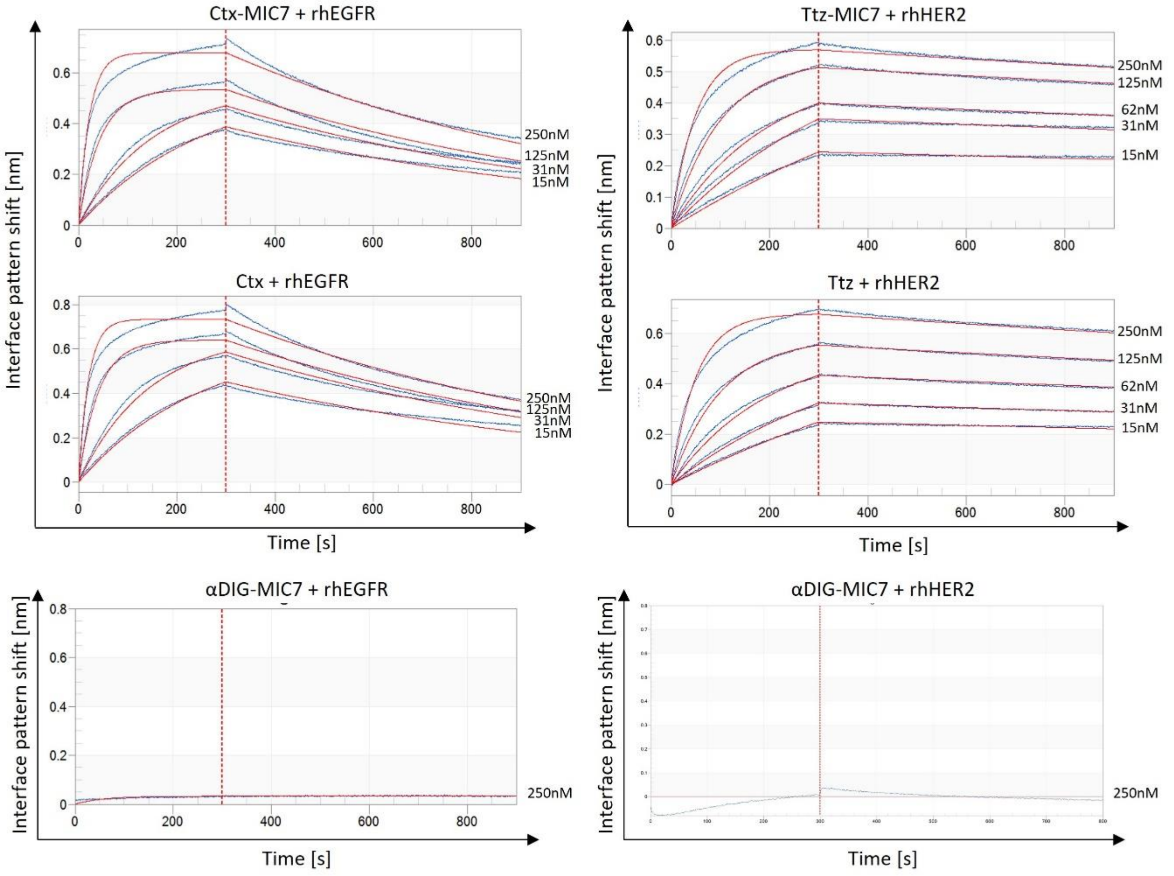


*Figure S 10 Binding analysis of parental antibodies and PROxAb shuttle conjugates with respective antigen via BLI. Association and dissociation were fitted by a 1:1 global full-fit binding model. The x-axis of the sensorgrams represents the wavelength interference shift in nm and the y-axis the time in seconds (s). Fittings are shown in red.*

*Table S 1 Biolayer interferometry analysis of recombinant human (rh) HER2 or EGFR binding. Respective dissociation constants ($K_D$) were calculated from fitting using FortéBio data analysis software. Curve fittings are included in the supplement (Figure S 10).*

| Antibody | Antigen | $K_D$ [M] |
|---|---|---|
| Ttz-MIC7 | rhHER2 | $2.2 \times 10^{-9}$ |
| Trastuzumab | | $2.5 \times 10^{-9}$ |
| Ctx-MIC7 | rhEGFR | $7.2 \times 10^{-9}$ |
| Cetuximab | | $6.2 \times 10^{-9}$ |

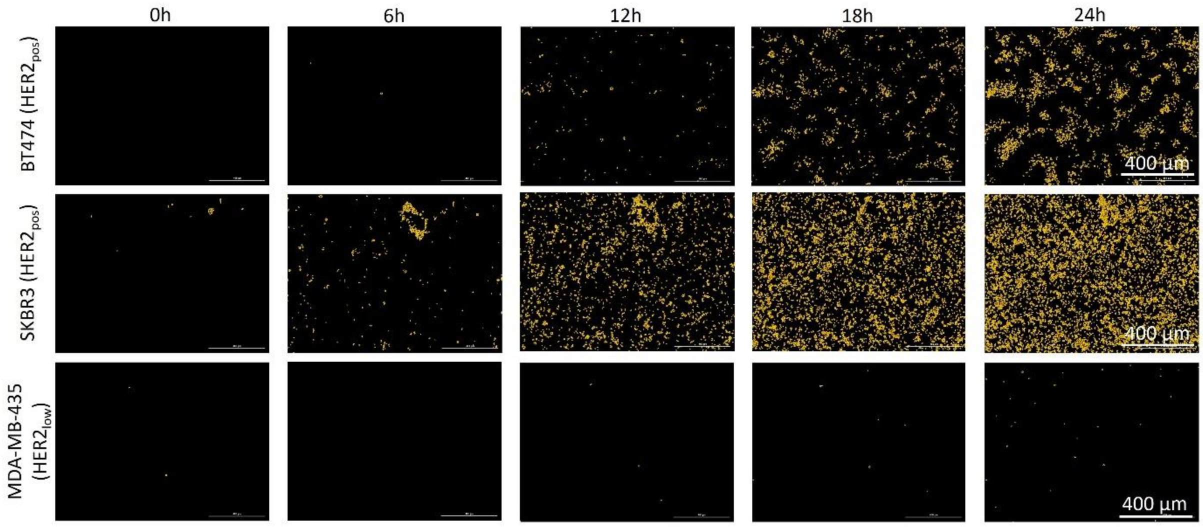


*Figure S 11 Live-cell imaging of HER2 positive and negative cell lines treated with pHAb-dye labeled Ttz-MIC7 for 24 hours. Images of Ttz-MIC7 internalization were selected at 0, 6, 12, 18 and 24 hours.*

*Table S 2 Binding affinities of conjugated VHH MIC7 to PROTACs GNE987, GNE987P and ARV771. *no binding detected up to a tested concentration of 0.5 µM. #determined off-rates are outside of instrument specifications and therefore be reported as $k_{off}$ < 0.0001 1/s; reported KD values are estimated by an assumption of $k_{on} > 1.0 \times 10^5$ [1/(M*s)]. $n.d. – not determined as $k_{off}$ is not measured unambiguously.*

| Antibody | PROTAC | $K_D$ [M] | $K_{on}$ [1/M*s] | $K_{off}$ [1/s] |
|---|---|---|---|---|
| Ttz-MIC7 | GNE987 | $< 1.0 \times 10^{-9}$ # | n.d. $ | $< 1.0 \times 10^{-4}$ # |
| | GNE987P | $3.82 \times 10^{-10}$ | $7.09 \times 10^{5}$ | $1.59 \times 10^{-4}$ |
| | ARV771 | $2.19 \times 10^{-9}$ | $1.83 \times 10^{5}$ | $4.19 \times 10^{-4}$ |
| Ctx-MIC7 | GNE987 | $< 1.0 \times 10^{-9}$ # | n.d. $ | $< 1.0 \times 10^{-4}$ # |
| | GNE987P | $3.83 \times 10^{-10}$ | $4.97 \times 10^{5}$ | $1.91 \times 10^{-4}$ |
| | ARV771 | $2.35 \times 10^{-9}$ | $2.29 \times 10^{5}$ | $5.04 \times 10^{-4}$ |
| Trastuzumab | GNE987, GNE987P, ARV771 | | no binding * | |
| Cetuximab | GNE987, GNE987P, ARV771 | | no binding * | |

*Table S 3 pHAb-dye conjugates generated in this study. Dye-to-antibody ratios were determined by photometric measurement using NanoDrop™ and LC-MS; means of both values are given.*

| Antibody | Dye-to-antibody ratio |
|---|---|
| Ttz-MIC7 | 6.7 |
| αDIG-MIC7 | 9.3 |
| Ttz | 6.5 |

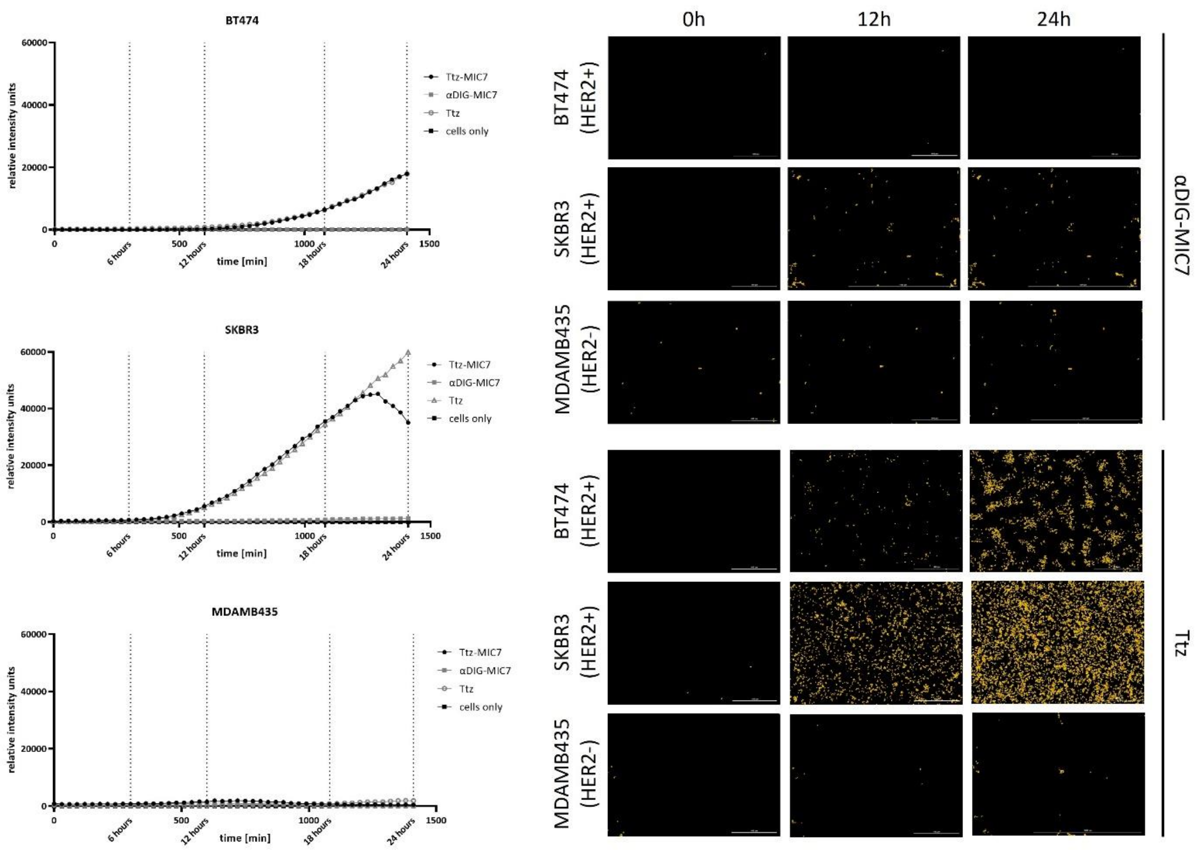


*Figure S 12 Internalization analysis of SKBR3, BT474 and MDA-MB-435 cancer cells incubated for 24 hours after treatment. Cells were treated with Ttz-MIC7, Ttz or αDIG-MIC7; untreated cells served as a reference. Relative intensity units were plotted for each cell line and treatment condition. Images (RFP channel: ex.: 531 nm, em: 539 nm and DAPI channel: ex.: 377 nm, em.: 447 nm) were taken with Cell Imaging Multimode Reader Cytation 5 (BioTek/Agilent) including 10x objective and LED intensity 10. An integration time of 14 ms and camera gain of 24 were used for the GFP channel, and 5 ms integration time and camera gain of 17 for the DAPI channel.*

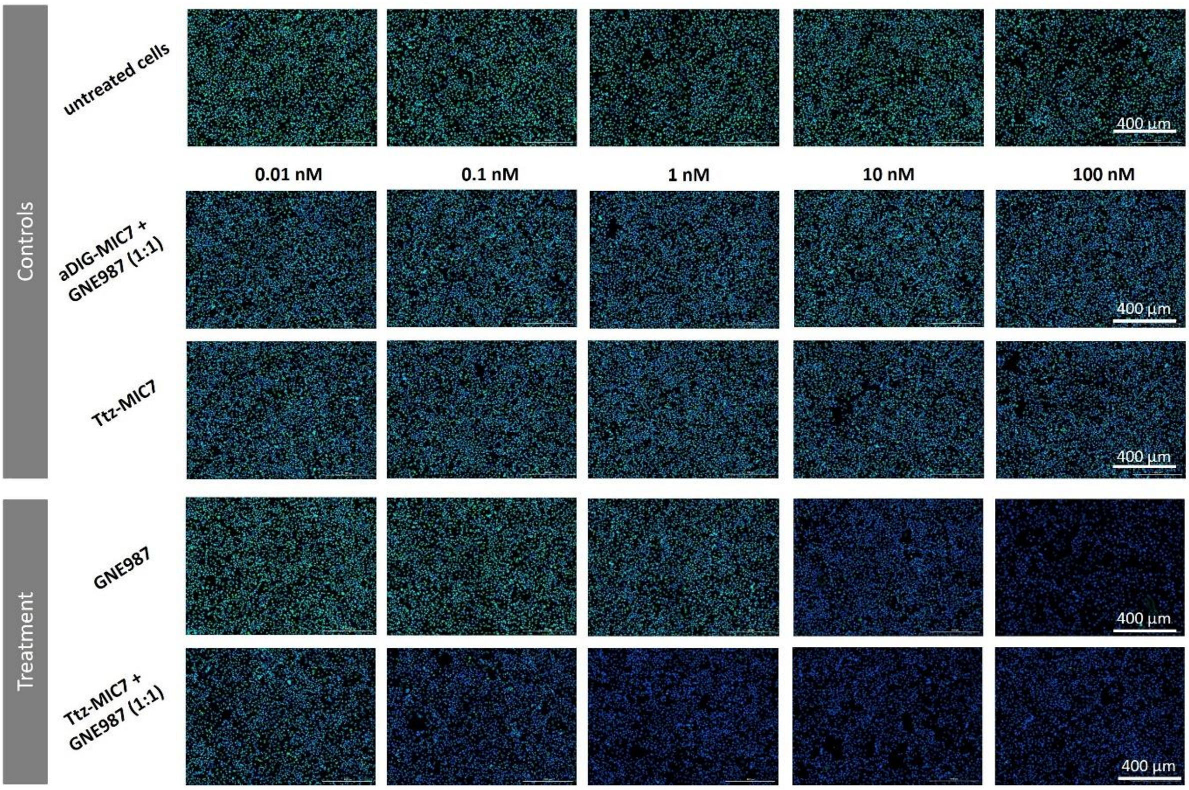


*Figure S 13 Immunofluorescence microscopy images of SKBR3 cancer cells incubated for 24 hours after treatment. Cells were treated with 10-fold serial dilution of GNE987, Ttz-MIC7 or αDIG-MIC7 loaded with GNE987, and unloaded Ttz-MIC7, while untreated cells were used as reference. Images (GFP channel: ex.: 469 nm, em: 525 nm and DAPI channel: ex.: 377 nm, em.: 447 nm) were taken with Cell Imaging Multimode Reader Cytation 5 (BioTek/Agilent) including 10x objective and LED intensity 10. An integration time of 48 ms and camera gain of 24 were used for the GFP channel, and 5 ms integration time and camera gain of 17 for the DAPI channel.*

*Table S 4 In vitro cytotoxicity data. $IC_{50}$ values of PROxAb shuttle-PROTAC combinations and respective controls derived from viability curves. nd: not determined, -: low or no impact on cell viability*

| $IC_{50}$ [M] | SKBR3 | | | BT474 | | |
|---|---|---|---|---|---|---|
| | GNE987 | GNE987P | *unloaded* | GNE987 | GNE987P | *unloaded* |
| Ttz-MIC7 | 4.2 x $10^{-10}$ | 5.5 x $10^{-10}$ | 1.5 x $10^{-9}$ | 6.7 x $10^{-10}$ | 7.3 x $10^{-10}$ | 1.1 x $10^{-9}$ |
| αDIG-MIC7 | 2.2 x $10^{-7}$ | >1.0 x $10^{-7}$ | >1.0 x $10^{-7}$ | >1.0 x $10^{-7}$ | 2.9 x $10^{-7}$ | >1.0 x $10^{-7}$ |
| Ttz | - | - | 8.5 x $10^{-10}$ | - | - | 8.0 x $10^{-10}$ |
| *free PROTAC* | 5.5 x $10^{-10}$ | 2.5 x $10^{-8}$ | - | 3.5 x $10^{-10}$ | 1.3 x $10^{-8}$ | - |
| | A431 | | | MDAMB468 | | |
| | GNE987 | GNE987P | *unloaded* | GNE987 | GNE987P | *unloaded* |
| Ctx-MIC7 | 1.8 x $10^{-10}$ | 4.3 x $10^{-10}$ | 3.2 x $10^{-9}$ | 1.8 x $10^{-10}$ | 2.4 x $10^{-10}$ | >1.0 x $10^{-7}$ |
| αDIG-MIC7 | nd | 3.0 x $10^{-8}$ | >1.0 x $10^{-7}$ | nd | >1.0 x $10^{-7}$ | >1.0 x $10^{-7}$ |
| Ctx | - | - | 3.1 x $10^{-9}$ | - | - | >1.0 x $10^{-7}$ |
| *free PROTAC* | 1.8 x $10^{-10}$ | 3.7 x $10^{-8}$ | - | 9.3 x $10^{-11}$ | 2.8 x $10^{-8}$ | - |
